# Agroforestry transition increases insect diversity and reorganizes soil function in a Mediterranean orchard

**DOI:** 10.64898/2026.05.28.728425

**Authors:** Solenn Patalano, Eleni Aplakidou, Evangelia Kagiali, Julia Miloczki, Maria Catana, Philippos Vardakas, Panagiotis Lembessis, Eleni Katsoni, Pantelis Hatzis, Giorgos A. Pavlopoulos, Sheila Darmos

**Affiliations:** Institute for Fundamental Biomedical Research (IFBR), BSRC ‘Alexander Fleming’, Vari 16672, Greece; Observatory of biodiversity dynamics, QUYFIN, Athens 17562, Greece; Division of Basic Sciences, University of Crete Medical School, Heraklion 71110, Greece; Climate Farmers, 12053 Berlin, Germany; The Southern Light Farm, Karyes 23067, Greece; Louca Distopia - Associação, 3780 Avelãs de Cima, Portugal; Department of Computational Biology, Mohamed bin Zayed University of Artificial Intelligence (MBZUAI), Building 1B, Masdar City, Abu Dhabi, United Arab Emirates

**Keywords:** Syntropic agroforestry, transects monitoring, soil metagenomics, sustainable farming systems

## Abstract

Agricultural intensification has reduced biodiversity and weakened ecosystem functions essential for sustainable crop production. Agroforestry has been proposed as a regenerative strategy to restore these functions through vegetation diversification, yet its ecological effects at the farm scale remain insufficiently documented, particularly in Mediterranean perennial systems. Here, adaptive field monitoring and soil shotgun metagenomics were combined to investigate ecological responses across a 10-year agroforestry transition gradient in a Mediterranean citrus and olive farm. Mature agroforestry plots supported higher insect richness and functional diversity while maintaining stable pollinator communities. Plant–insect interaction analyses suggested that potential pest activity was largely associated with spontaneous and supportive vegetation rather than crops. In contrast, soil communities showed limited changes in overall richness but substantial compositional and functional restructuring, including enrichment of the arbuscular mycorrhizal fungus *Rhizophagus* and proteins with nutrient-cycling functions in advanced plots. Together, these findings suggest that agroforestry transition can promote functional diversification above and below ground and highlight the value of farmer-led regenerative transitions coupled with integrative ecological monitoring for the development of more resilient agricultural systems.

## Introduction

By simplifying habitats and reducing biodiversity, agricultural intensification has progressively disconnected crop production from the ecological interactions that sustain it, including pollination, pest regulation, and nutrient cycling, increasing dependence on external inputs while undermining long-term ecosystem stability and resilience (Power, 2010). These losses challenge the long-term sustainability and resilience of agri-food systems, as biodiversity is a key driver of ecosystem functioning and stability (Cardinale et al., 2012). In response, regenerative agriculture aims to reconcile productivity with ecological functioning by promoting practices that enhance biodiversity while maintaining or improving crop yields (Altieri, 1999; Wezel et al., 2014).

Agroforestry has emerged as a key diversification strategy within this transition, by intentionally integrating trees, supportive plant species, and spontaneous vegetation into crop production system to enhance ecosystem functioning (Atangana et al., 2014; Smith et al., 2012). Through the combination of tree canopies, cultivated plants, and spontaneous ground cover, agroforestry systems increase habitat stratification and plant diversity, creating a wider range of ecological niches and resources within agricultural landscapes. These more complex environments are expected to support a broader diversity of organisms and ecological interactions while moderating fluctuations in local environmental conditions. (Elevitch et al., 2018; Sollen-Norrlin et al., 2020). Increased plant and flower diversity may enhance pollinator and natural enemy communities by providing additional food resources and habitats, while diversified organic inputs from vegetation may influence soil microbial community composition and functioning. However, despite strong theoretical support, empirical evidence linking agroforestry transition to functional ecological outcomes at the farm scale remains limited, particularly in perennial Mediterranean systems (Altieri et al., 2024; Mosquera-Losada et al., 2023).

Insect communities are central to agroecosystem functioning, as they mediate both ecosystem services and disservices across trophic levels. Pollinating insects are essential for the productivity and quality of many crops (Klein et al., 2007; Potts et al., 2010), and global syntheses have shown that both insect biodiversity and non-bee pollinators substantially contribute to crop production and ecosystem resilience (Dainese et al., 2019; Rader et al., 2015). Conversely, insect pests remain a major source of crop damage and yield loss worldwide, a pressure expected to intensify under climate change through increased pest growth, consumption, and geographic expansion, particularly in simplified agricultural systems with reduced ecological regulation (Ma et al., 2025; Tonnang et al., 2022). However other insects contribute to biological pest control through predation and parasitism, while plant diversification and landscape complexity have been shown to enhance natural enemy communities and reduce pest pressure in agricultural systems (Jonsson et al., 2015; Letourneau et al., 2011). Analyses of plant–pollinator interactions further suggest that increasing plant diversity can promote pollinator richness and strengthen ecological interaction networks within agroecological farms (Astegiano et al., 2024). Despite this complexity, monitoring of insect diversity and plant–insect interactions in agricultural contexts remains largely centred on pollinators and is often based on single-time assessments that fail to capture temporal dynamics and shifts in functional diversity. As a result, existing approaches rarely provide farmers with accessible monitoring tools capable of detecting broader ecological trends, seasonal pest emergence, or changes in ecosystem functioning at the farm scale.

Since vegetation diversification not only modifies habitat heterogeneity and floral resources above ground, but also organic inputs below ground, agroforestry transitions are expected to influence soil biodiversity and microbial functioning. Indeed, soil biodiversity plays a fundamental role in sustaining agroecosystem functions through organic matter decomposition, nutrient cycling, soil structure maintenance, and plant–microbe interactions (Xiong & Lu, 2022). Agroforestry is expected to influence these below-ground communities through changes in vegetation composition, litter inputs, root exudation, and microenvironmental conditions, potentially reshaping microbial assembly and ecosystem functioning (Jing et al., 2022; Vaupel et al., 2025). However, soil community responses to agroforestry transition remain relatively understudied. Existing studies suggest that soil microbial communities can respond predictably to climate events, agricultural management, and restoration practices, making them useful bioindicators of ecosystem response, soil health, and recovery (Armbruster et al., 2021; Knight et al., 2024; Sánchez-Cueto et al., 2025). While conventional soil assessments often rely on physicochemical indicators or targeted molecular approaches, recent advances in shotgun metagenomics provide complementary opportunities to characterize multitrophic diversity alongside functional gene repertoires across agroecosystems (Clark et al., 2021; Fierer, 2017).

In this study, adaptive transect-based field monitoring and soil shotgun metagenomics were combined to investigate above- and below-ground species dynamics at Southern Lights Farm, a Mediterranean orchard undergoing a gradual transition to agroforestry. We hypothesized that increasing vegetation diversification and habitat complexity across the agroforestry transition would promote insect diversity and beneficial ecological functions above ground while also reshaping soil microbial taxonomic and functional composition below ground. This study aimed to (i) quantify plant and insect community dynamics, including taxonomic composition, richness, abundance, and functional roles, across a farm-scale agroforestry gradient; (ii) assess how vegetation diversification influences insect-mediated ecosystem services and pest dynamics; and (iii) characterize soil taxonomic composition and functional potential to determine whether agroforestry transition is associated with the emergence of a below-ground ecological signature. By integrating these analyses within a single farm system, this work provides empirical evidence on the ecological effects of agroforestry systems that can inform management-relevant strategies for sustainable agricultural transitions.

## Results

### Farm structure and agroforestry planting system

The Southern Lights Farm is organised into five adjacent plots (A–E) differing in their year of transition to agroforestry and forming a gradient of system maturity across the farm (**Figure 1A, Supplementary Fig. S1**). To characterise plant community structure, scalable transects were designed to be proportional to plot size, representing on average approximately 2% of each plot area, with a mean length of 39.9 ± 7.0 m (mean ± SD across all plots and surveys). All visible plant species were recorded monthly during the plant community survey conducted between March and June 2024 (see M&M).

**Figure 1.**
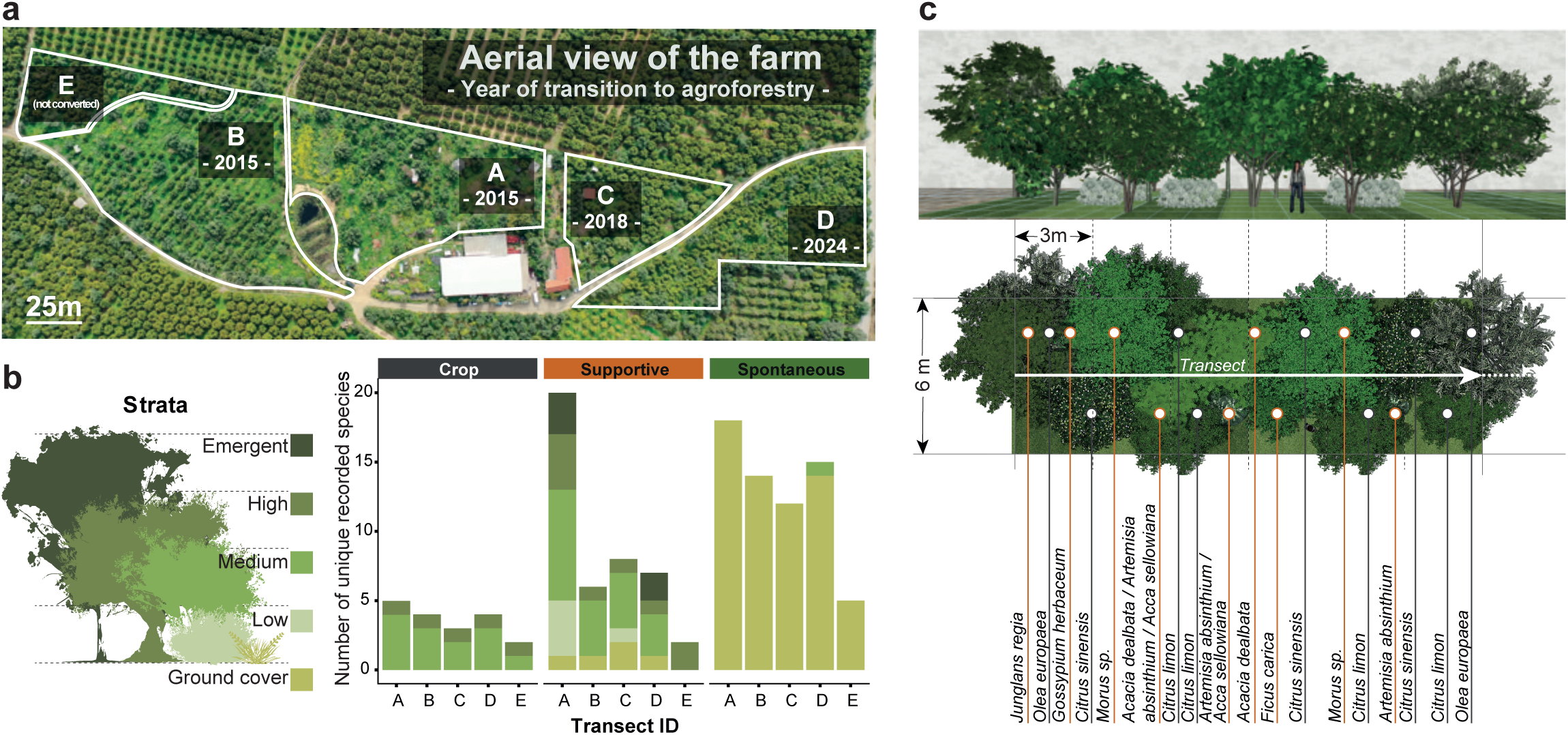
Farm structure and agroforestry planting system composition. **(a)** Aerial view of the Southern Lights Farm showing the spatial arrangement of the five plots (A–E) and year of transition to agroforestry. Plot boundaries are delineated in white. **(b)** Visual representation of the agroforestry vertical strata used for plant classification (left) and bar plots showing the number of unique plant species recorded per transect, grouped by functional role and stacked by strata (right). Values reflect planting composition rather than effort-standardised richness and are derived from the plant community survey conducted between March and June 2024 for each transect. Mean transect length (± SD across four monthly surveys): A, 43.0 ± 2.4 m; B, 35.3 ± 2.9 m; C, 48.5 ± 1.0 m; D, 42.5 ± 2.6 m; and E, 28.5 ± 1.7 m. **(c)** Example of a transect section (0–18 m) from plot A illustrating the agroforestry planting system as monitored in Spring 2024. Panels present a side view of the agroforestry layout (upper), a bird’s-eye view of the same section (middle), and the corresponding plant species identities (bottom). Crop species (dark grey lines) and supportive species (orange lines) are indicated, while all spontaneous species of this section (*Lamium bifidum, Geranium molle, Vicia sativa, Galium aparine, Parietaria judaica, Taraxacum officinale, Medicago polymorpha, Bidens bipinnata, Oxalis pes-caprae, Ranunculus repens, Sinapis arvensis, Urtica dioica, Lysimachia foemina, Lysimachia foemina v. white, Stellaria media, Fumaria capreolata, Fumaria sp., Anthriscus sp., Malva sylvestris, Convolvulus sp., Solanum nigrum*) were virtually represented as a green ground layer. Visuals were performed using Realtime Landscaping Architect (RLA), Version 2020. The transect monitoring corridor between two tree lines is indicated in white.

In total, 56 plant species were recorded along the five transects (**Table S1**). Crop diversity was limited to six species, comprising olive (*Olea europaea)* and five citrus species, representing the high and medium canopy strata, respectively. Crop species composition exhibited a uniform distribution across all transects, reflecting the shared orchard backbone and consistent management of the main cultivated trees across parcels (**Figure 1B**). In contrast, supportive plant species showed strong heterogeneity between transects. A total of 25 support species were recorded, distributed across all five vertical strata and reflecting deliberate planting to build agroforestry structure. Transect A showed a substantially higher number of support species than all other transects (**Figure 1C**). In comparison, transects B, C, and D shared a more limited and partially overlapping set of supportive woody species, mainly comprising fruit-yielding species such as *Morus sp*., *Prunus spp.*, *Punica granatum*, and, in the most recent conversion, fast-growing biomass trees including *Eucalyptus globulus* and *Populus alba*. Herbaceous support species (*Vicia sativa*, *Medicago polymorpha*, *Bidens bipinnata)* were consistently recorded in all transects and contributed primarily to the ground cover and the medium strata. A further 25 spontaneous plant species were recorded, with broadly similar composition among transects, except for transect E, which showed a reduced set of spontaneous species. A core group of spontaneous species was consistently recorded across all transects, comprising *Sinapis arvensis*, *Oxalis pes-caprae*, *Taraxacum officinale*, *Malva sylvestris*, and *Parietaria judaica*. Throughout the farm, spontaneous species were associated almost exclusively with the ground-cover stratum. Together, these patterns indicate that the transects captured both a shared baseline spontaneous vegetation and plot-specific structural differences associated with agroforestry establishment and maturity.

### Agroforestry transition is associated with increased insect richness, abundance, and functional diversity

To assess how the agroforestry transition influences insect functional diversity and associated ecosystem services, we examined seasonal variation in floral resources and insect community dynamics across the farm. This analysis included not only pollinators but also insect groups with ecological roles relevant to crop production, including natural predators, potential pests, and taxa with mixed functional roles across life stages. Insect communities were quantified across 23 transect surveys conducted during the insect visitation surveys across all five plots during spring 2024 (mean survey duration 23.7 ± 13.2 min). Surveys were performed under comparable abiotic conditions (mean temperature 28.8 ± 3.9 °C; wind speed < 3 m·s⁻¹).

A total of 814 flowering plant observations were recorded, corresponding to 47 unique flowering plant species. All main crop species were flowering during the survey period, together with 15 supportive species, including *Acacia dealbata, Rosmarinus officinalis, Punica granatum, and Prunus spp.* The majority of flowering taxa (25 species) consisted of spontaneous herbaceous plants, including *Lysimachia spp., Malva sylvestris, Oxalis spp., Taraxacum officinale, and Parietaria judaic*a. In parallel, 300 insect observations were recorded and assigned in the field to 20 operational morphogroups. Of these, 70% were further identified taxonomically, allowing classification into 34 families and 43 species (**Table S2**). Beetles (Coleoptera) represented the most taxonomically diverse group (8 families, 10 species). Several families of wild pollinator bees were recorded, including *Xylocopa violacea* (Apidae), *Andrena sp.* (Andrenidae), and *Lasioglossum spp.* (Halictidae), together with hoverflies (*Episyrphus balteatus*, Syrphidae) and butterflies (*Vanessa spp*., Nymphalidae). Taxa with potential negative impacts on crops were also observed, including sap-sucking Hemiptera (*Beosus quadripunctatus, Cydnus aterrimus, Closterotomus spp., Graphosoma italicum*), leaf-feeding insects (*Pieris rapae, Anacridium aegyptium*), and rarer species such as the Greek antlion *Nemoptera coa*, the cuckoo wasp *Holopyga fervida*, and the moth *Acontia trabealis*.

Flowering plant species richness declined consistently from early to late spring across all transects (**Supplementary Fig. S2**), reflecting seasonal phenology and progressive reduction of floral resources. Insect species richness followed a similar overall trend; however, insect Shannon diversity peaked during mid-spring (April–May), indicating a transient diversification of insect communities during the period of floral availability. Lower Shannon diversity values observed in March coincided with honeybee dominance, likely reflecting the presence of transhumant hives during peak citrus flowering in the surrounding landscape. The unconverted parcel (transect E) consistently exhibited the lowest insect diversity throughout the monitoring period, whereas transects representing more advanced stages of agroforestry transition — particularly transect A — maintained higher and more variable insect diversity. By June, flowering plant availability was largely restricted to a small subset of spontaneous herbaceous species, primarily *Parietaria judaica* and *Convolvulus* spp., coinciding with a marked decline in insect diversity. Across transects and months, insect species richness increased significantly with flowering plant species richness (linear mixed-effects model; β = 0.47 ± 0.10 SE, t = 4.67, p = 0.00084; **Figure 2a**), indicating a consistent positive association between local floral diversity and insect diversity across the agroforestry transition gradient.

**Figure 2.**
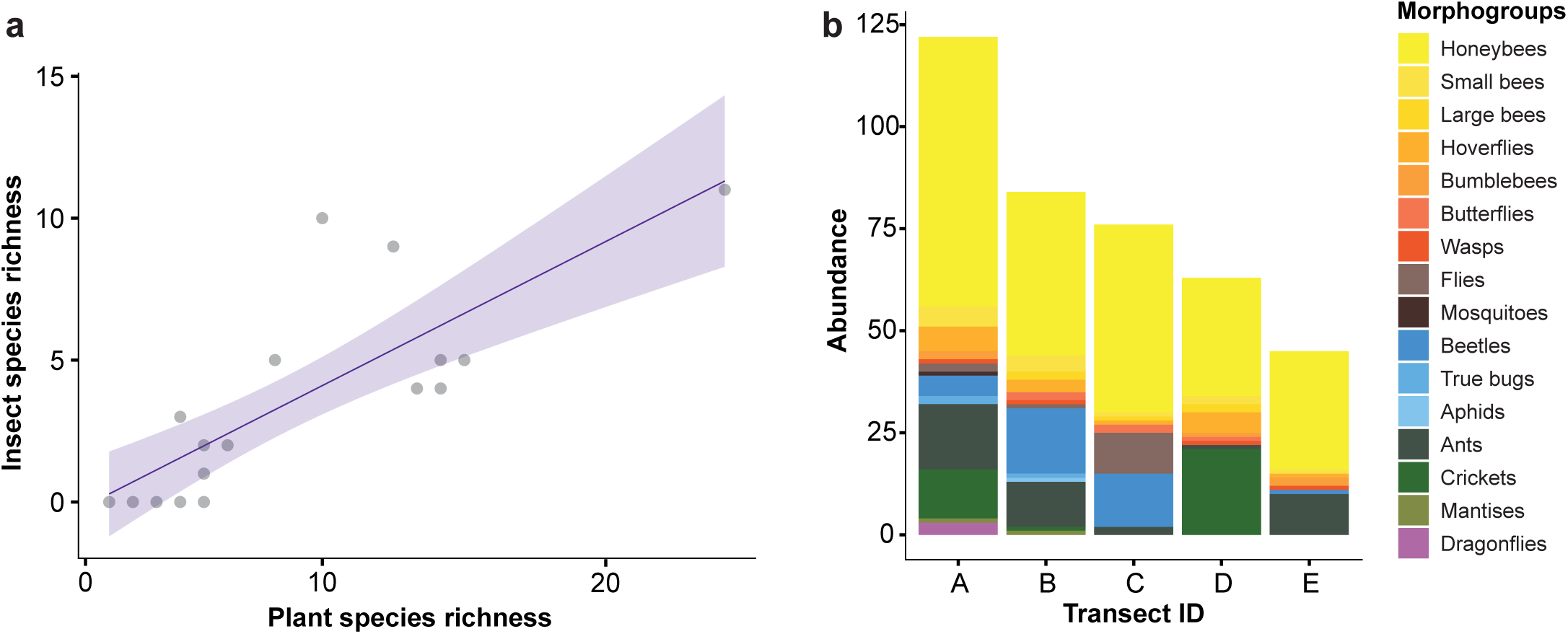
Plant diversity shapes insect richness and community composition. (a) Scatter plot showing the relationship between flowering plant species richness and insect species richness. Points represent individual transect–month observations. The purple line shows the overall linear trend and the shaded area represents the 95% confidence interval. (b) Stacked bar plots showing the cumulative abundance of insect morphogroups pooled across all Spring samples for each transect. Abundances are based on frequency-weighted observations from the first 27 m of each transect.

To determine whether increased insect richness reflected changes in community composition, we examined morphogroup-level abundance patterns across transects (**Figure 2b**). While dominant pollinator groups such as honeybees were consistently present across all plots, compositional differences among transects were primarily driven by the addition of new insect morphogroups rather than by replacement of existing ones; assemblages in the unconverted parcel (transect E) largely represented a subset of those observed in more advanced agroforestry stages. Specifically, agroforestry plots exhibited the emergence of beetles (Coleoptera), true bugs (Hemiptera), aphids, and orthopterans (e.g. crickets and grasshoppers). Collectively, these patterns indicate that agroforestry transition is associated with increased insect richness and functional heterogeneity, translating plant diversification into more complex above-ground insect communities.

### Agroforestry vegetation enables pollination services while buffering pest pressure

Since greater insect diversity does not necessarily imply beneficial effects on agricultural production, we then examined the distribution of functional groups of insects interacting with flowering vegetation across the farm. A total of 235 interactions between flowering plants and insects were observed (**Figure 3a**). The largest proportion of unique plant–insect interactions was observed on spontaneous vegetation (44.3%), followed by crop plants (35.3%), and supporting plant species (20.4%). It should be noted that several flowering support species were never observed interacting with insects during the monitoring period, including *Acca sellowiana, Tithonia diversifolia, Tradescantia pallida,* and *Erythrostemon gilliesii*. Their tropical or subtropical origins may partly explain their limited interaction with local insect communities under Mediterranean spring conditions, although limited sampling cannot be excluded.

**Figure 3.**
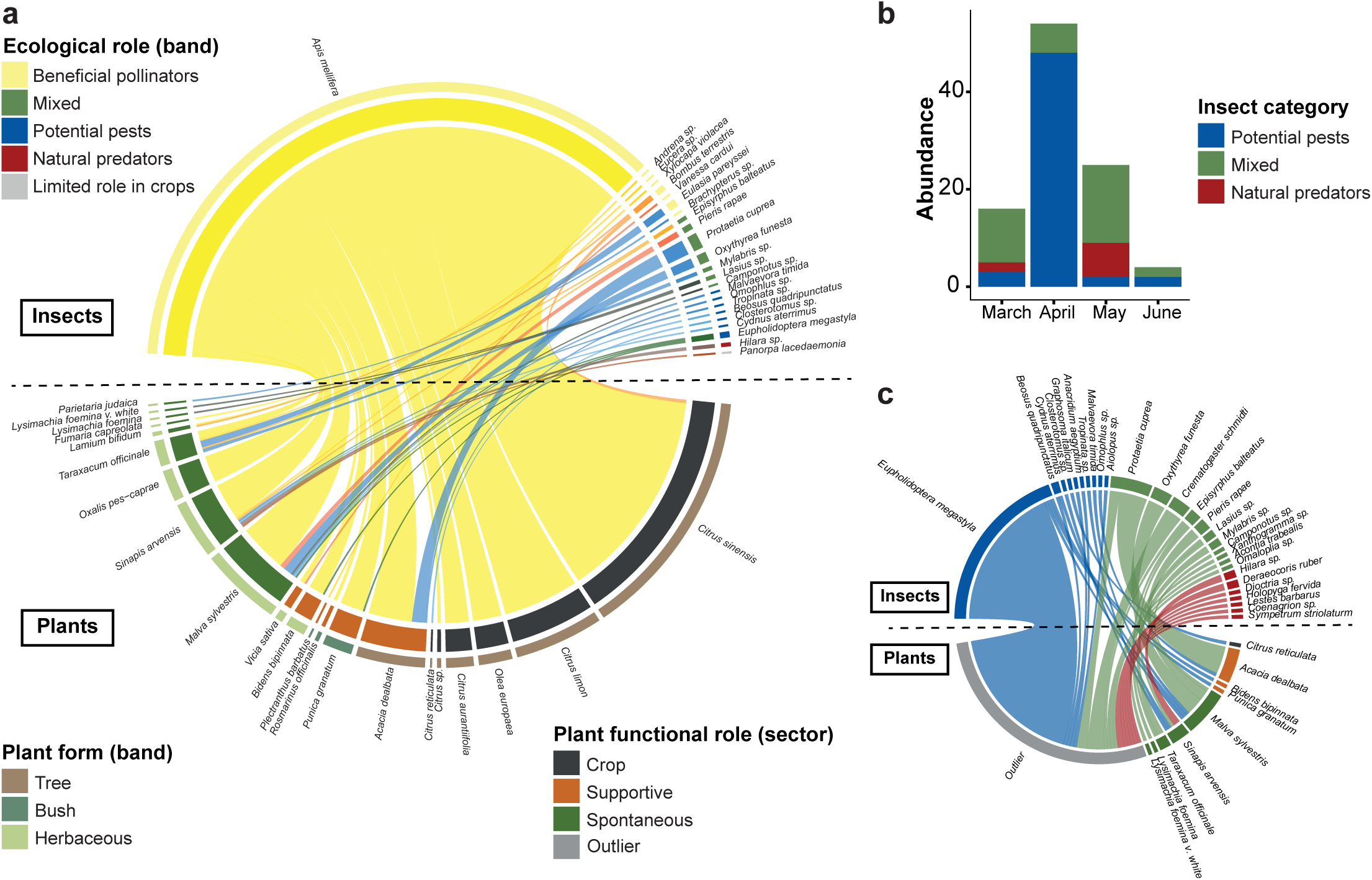
Supportive and spontaneous vegetation shape insect interactions and buffer pest pressure. (a) Chord diagram showing frequency-weighted plant–insect interaction network at the farm scale between flowering plant species (lower sectors, coloured by plant functional role) and insect species (upper sectors, coloured by ecological role), pooled across all transects and sampling months. Link widths are proportional to summed interaction frequency. Outer bands indicate plant growth form (plants) and ecological role (insects). (b) Bar plots showing monthly distribution of non-pollinator insect abundance categorised by ecological role and pooled across transects. Abundances are based on frequency-weighted observations. (c) Chord diagram showing frequency-weighted associations between non-pollinator insect species (upper sectors, coloured by ecological role) and plant species (lower sectors, coloured by plant functional type). Outliers correspond to observations of insects present on vegetation or within transect sections without direct contact with floral or reproductive plant structures. Link widths are proportional to the summed observation frequency.

Interaction types were highly uneven across insect taxa. Honeybees (*Apis mellifera*) alone accounted for 55.4% of all recorded interactions, while the inclusion of butterflies, hoverflies, and other bees, increased the total proportion of pollinator interactions to 69.6%. The single most frequent interaction involved *Citrus sinensis* and *A. mellifera*, representing 27.3% of all recorded interactions. Notably, 98.6% of insect interactions involving crops were attributable to beneficial pollinators, with only one interaction recorded with a potential pest species (*Closterotomus sp.*). Interactions involving other insect functional groups were comparatively rare and were predominantly associated with non-crop vegetation. Potential pest taxa (*Beosus quadripunctatus, Cydnus aterrimus, Eupholidoptera megastyla, Malvaevora timida, Tropinata sp., Omophlus sp.*) interacted primarily with spontaneous plants (55.6%) and supporting species (33.3%). These patterns indicate that, despite the emergence of additional insect functional groups during the agroforestry transition, crop plants remained predominantly visited by beneficial pollinators, while potential pest activity was largely confined to spontaneous and supporting vegetation.

Abundance analyses further revealed pronounced temporal and spatial structuring of non-pollinating insect activity during the spring sampling period. A marked peak in abundance was observed in April, driven primarily by *Eupholidoptera megastyla*, a bush-cricket not typically classified as an agricultural pest but whose omnivorous feeding behaviour and local densities may cause foliar damage (**Figure 3b**). This peak was followed by a sharp decline in May. In contrast, natural predator activity increased later in the season, particularly in May, and was mainly driven by predatory odonates and dipterans (e.g. *Coenagrion sp., Lestes barbarus, Sympetrum sp., Dioctria sp.*). Importantly, potential pest activity was largely confined away from flowering crops (**Figure 3c**). Overall, 65.7% of pest individuals were recorded as outlier observations not interacting with flowering parts, while the remaining individuals were associated with supporting (11.1%) and spontaneous vegetation (22.2%). Although replication is limited and abundance estimates are semi-quantitative, the temporal and spatial trends observed here are consistent with a buffering role of spontaneous and supporting vegetation in agroforestry systems, reducing pest pressure on crops while facilitating seasonal predator build-up.

### Soil environment and taxonomic composition is highly homogeneous across the farm

To evaluate soil characteristics across the farm, we combined physicochemical and metagenomic analysis of soil samples along the established transects. Soil surface pH averaged 6.45 ± 0.70 across all monitoring, with no significant differences detected across transects (n=38, Kruskal–Wallis test, p = 0.054). This homogeneous pH suggests similar soil chemical conditions across parcels, with no detectable differences linked to water or input management. For soil biodiversity assessment, DNA extracted from 5 composite soil samples were subjected to untargeted high-throughput metagenomic sequencing. The sequencing generated a total of 6,646,089 reads distributed across 20 libraries (5 soil samples x 4 technical replicates), with the size of each library ranging from 104,902 to 587,887 reads, providing sufficient and comparable sequencing depth to capture the majority of detectable soil taxonomic diversity (**Supplementary Fig. S3**).

At the farm level (across all soil samples), most reads were assigned to bacteria (95.2%), followed by archaea (2.7%), eukaryotes (2.1%), and viruses (<0.1%) within the proportions typically recovered in bulk soil metagenomes (**Figure 4a**). Within Bacteria, the community was dominated by the phylum Actinomycetota, which accounted for 60.2% of bacterial reads, followed by Pseudomonadota (23.2%) and Bacillota (7.3%). In total, 825 bacterial genera were identified. Archaeal reads were mostly assigned to Nitrososphaerota (90.3%), with *Nitrosocosmicus oleophilus* representing the most abundant predicted species within this phylum. Eukaryotic reads showed a more even distribution across kingdoms (**Figure 4b**): Fungal DNA represented 38.1% of eukaryotic reads encompassing 25 genera (**Table S3**), while plant DNA accounted for 37.3% with 29 genera identified (**Table S4**) and animal-derived DNA represented 22.6% of eukaryotic reads with 35 genera identified (**Table S5**). Viral reads were predominantly assigned to *Caudoviricetes spp.*, in line with previous analysis showing the dominance of this tailed bacteriophage in soil ecosystems (Bi et al., 2021).

**Figure 4.**
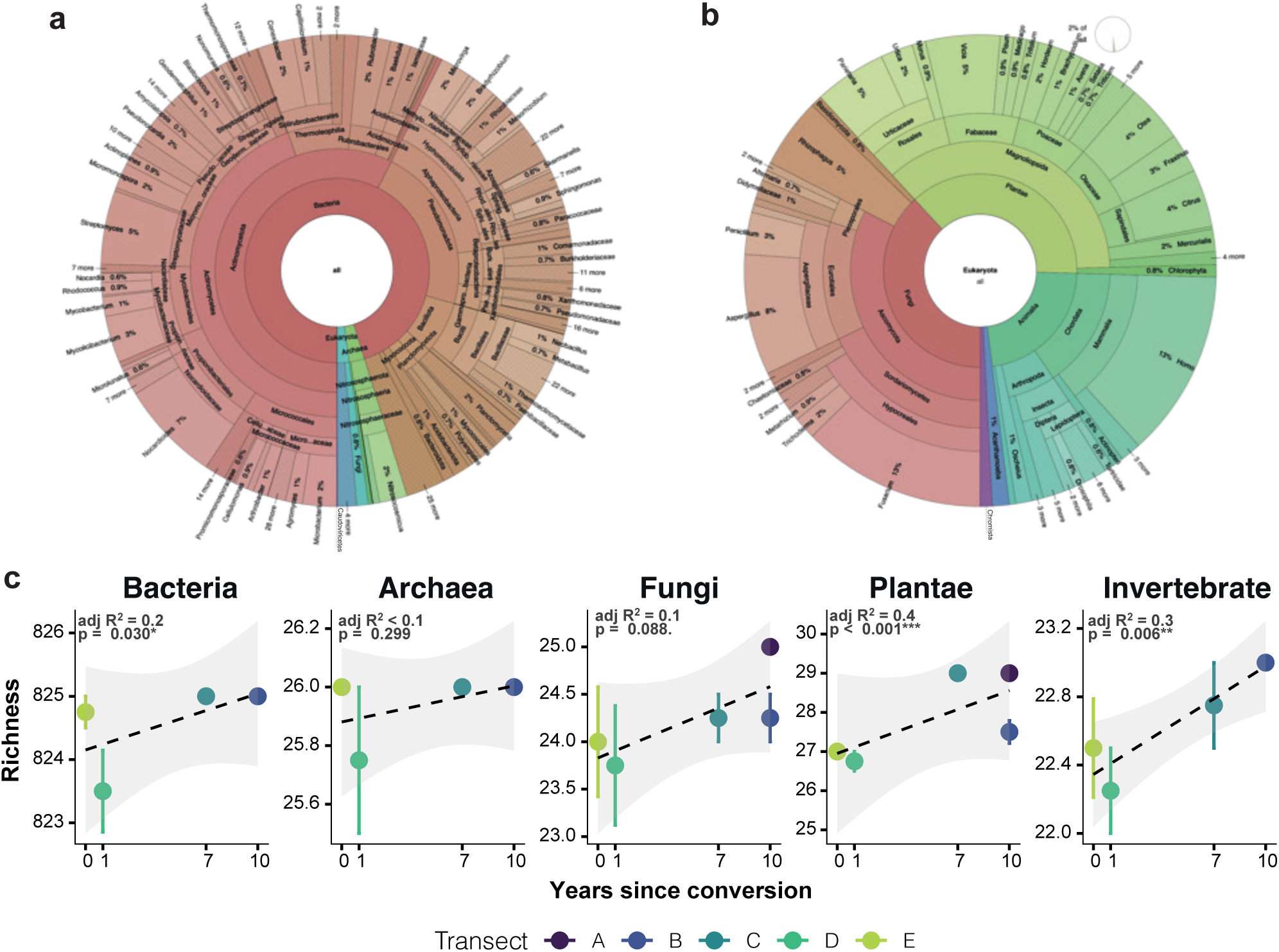
Taxonomic composition of soil communities across the farm. Krona visualization of the relative abundance of all classified taxa across all soil samples (a) and a zoomed-in view of the Eukaryota domain (b), displayed from Domain to Genus level. Taxa were agglomerated to the genus level and filtered to retain only genera reaching a relative abundance of at least 0.01% in at least 15% of samples across the dataset (n = 1,046 genera). Krona plots were generated using KronaTools for hierarchical visualization of metagenomic data (Ondov et al., 2011). (c) Scatter plot showing the relationship between mean genus richness and years since agroforestry conversion for major taxonomic groups. Points represent mean genus richness ± standard error (SE) calculated across technical replicates within each transect. Richness was calculated at the genus level for Bacteria and Archaea Kingdoms, for Fungi and Plantae Phyla and for Invertebrates by grouping genera belonging to the phyla Arthropoda, Nematoda, and Mollusca. Dashed lines show linear regression trends across transects, with shaded areas indicating 95% confidence intervals. Adjusted R² values and p-values from linear models are shown for each group.

Genus-level richness was broadly conserved across transects for all major taxonomic groups (**Figure 4c**). Mean genus richness varied modestly along the agroforestry gradient, with weak positive trends observed for Bacteria, Plantae, and Invertebrates, and no significant trend detected for Archaea or Fungi. Across all groups, effect sizes were small, corresponding to differences of approximately 2–3 genera over a 10-year conversion period, indicating limited variation in the total number of detected genera. Genus prevalence analysis further revealed a high degree of taxonomic overlap among transects, with 99.8% of genera detected in all five transects. Only two plant genera showed transect specific absences: *Vicia sativa* was undetected in transects B and E, and *Pisum sativum* from transect E. No genera were uniquely associated with a single transect. Consistently, negligible taxonomic turnover was observed across transects, as indicated by very low Jaccard dissimilarities (range: 0–0.002). Taken together, these results indicate a highly homogeneous soil taxonomic composition across the farm, with near-complete conservation of genus-level identity among transects.

### Agroforestry transition drives plot-specific community restructuring with a small emerging agroforestry signature

Community structural dynamics were then examined to determine whether agroforestry transition was associated with temporal changes in soil taxonomic organisation.

Across the 10-year conversion period, alpha-diversity responses differed strongly among biological groups (**Supplementary Fig. S4**). Plant communities showed a consistent and significant increase in Shannon diversity, Simpson diversity, and evenness with years since conversion, indicating diversification through reduced dominance and more even coexistence of taxa. This directional shift reveals a structural reorganisation of plant communities despite minimal taxonomic turnover and provides a below-ground confirmation of the effect of agroforestry establishment. By contrast, bacterial communities exhibited a modest but significant decline in Shannon diversity and evenness during agroforestry transition, highlighting the increased dominance of a subset of bacterial genera, whereas archaeal and fungal diversity indices showed no significant temporal trend despite evidence of temporal compositional rearrangements. Interestingly, invertebrate diversity declined significantly over time across all indices, reflecting increasing dominance within the invertebrate assemblage; this pattern appears largely driven by the expansion of an unidentified nematode-related taxon in transects A and B (See below and **Tables S5, S6**). Taken together, these results do not support a gradual successional trajectory with time since conversion, but instead point toward a heterogeneous, plot-specific restructuring of the non-plant communities.

To determine whether this restructuring was also reflected in community composition, beta-diversity patterns were examined among transects. Principal Coordinates Analysis (PCoA) based on Bray–Curtis community dissimilarities positioned the centroids of the converted plots (A, B, C, D) multidirectionally and nonlinearly in ordination space relative to the unconverted reference plot (E) (**Figure 5a**). Divergence from the reference followed an increasing deviation in composition in the order E < C < B < A < D (Bray–Curtis dissimilarities E: 0.0187 ± 0.0003; C: 0.118 ± 0.0005; B: 0.131 ± 0.0008; A: 0.170 ± 0.0011; D: 0.195 ± 0.0023), a pattern that does not align with a gradient based on years since conversion. Moreover, pairwise comparisons of Bray–Curtis dissimilarity among all transects revealed that plant communities exhibited the greatest compositional divergence (range 0.445–0.797), followed by fungal (0.198–0.480) and invertebrate communities (0.177–0.476), whereas bacterial (0.115–0.227) and archaeal communities (0.033–0.193) showed moderate variation (**Figure 5b**). Interestingly, the largest dissimilarities were observed between transects C and D for plants and fungi, and between transects A and D for invertebrates, reinforcing the idea that community restructuring was heterogeneous and strongly plot-specific rather than temporally ordered along the agroforestry conversion gradient.

**Figure 5.**
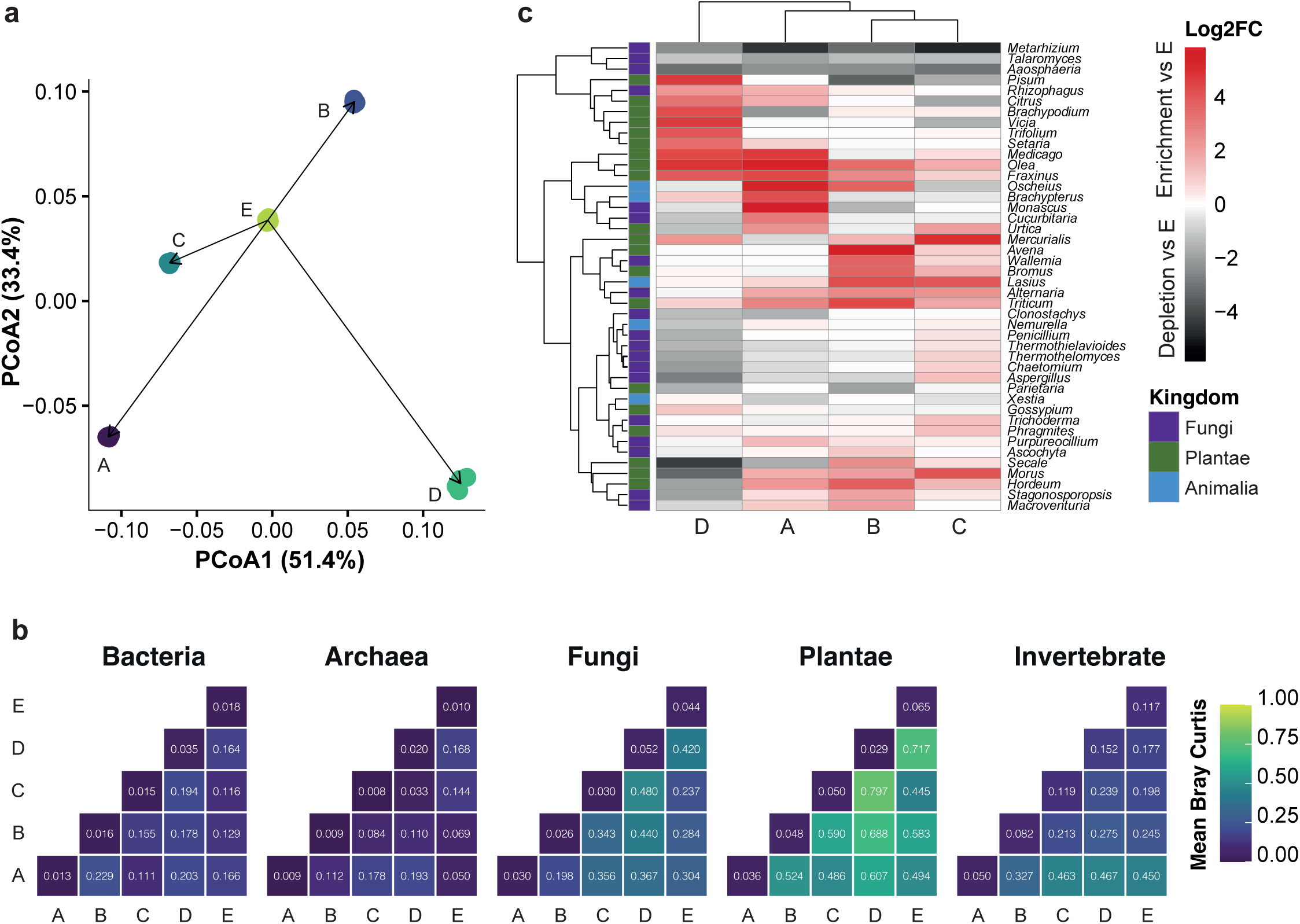
Community compositional divergence and taxa driving agroforestry transition. (a) Principal Coordinates Analysis (PCoA) based on Bray–Curtis dissimilarities calculated from relative abundance profiles at the genus level. Each dot represents a soil library coloured by transect origin. Arrows originating from the centroid of the unconverted transect (E) indicate the direction and magnitude of compositional displacement of each converted transect centroid. Percent variance explained by each axis is shown. (b) Mean Bray–Curtis dissimilarity among transects calculated from relative abundance data. Values represent the mean pairwise dissimilarity between all libraries belonging to each transect pair. Diagonal cells indicate within-transect dissimilarity (technical baseline). Warmer colours indicate greater compositional divergence. (c) Heatmap showing the log₂ fold change (log2FC) of the 38 genera that were significantly differentially abundant relative to the reference transect (E) based on the ANCOM-BC2 analysis (q < 0.05 and |log2FC| ≥ 1). Columns represent the agroforestry transition transects (A–D), and rows represent genera detected in the soil metagenomic dataset. Positive values (red) indicate enrichment relative to the reference transect, whereas negative values (black) indicate depletion. Genera are hierarchically clustered based on similarity in their differential abundance profiles across transects. Row annotations indicate the taxonomic kingdom of each genus (Fungi, Plantae, or Animalia).

To identify the taxa responsible for the compositional divergence among transects, differential abundance analysis was performed on fungal, plant, and invertebrate genera using ANCOM-BC2. Across the converted transects, 38 genera showed significant differential abundance relative to the unconverted reference transect E (FDR-adjusted q < 0.05 and ≥ 2-fold change; **Table S6**, **Figure 5c**). As expected, the number of significantly different genera varied among plots: 19 genera in transect A, 15 in transects B and C, and 23 in transect D, confirming that agroforestry transition is associated with taxonomic restructuring of varying magnitude, with transects D and A showing the most significative divergence.

Seven genera were identified as the main contributors to the multidirectional taxonomic restructuring and plot-specific responses associated with agroforestry transition. *Citrus* was significantly enriched in transects A and D but significantly depleted in C, likely reflecting variability in planting practices across the farm. Other plants showed similar significant contrasting patterns, including *Hordeum* (enriched in A, B, and C but depleted in D), the nettle *Urtica* (enriched in A and C but depleted in D) and an herbaceous species belonging to the genus *Brachypodium* (enriched in D but depleted in A). The saprotroph and foliar endophytic fungi *Aspergillus* (enriched in C but depleted in D) and the plant-pathogenic fungus *Stagonosporopsis* (enriched in B but depleted in D) were also found. Finally, a nematode-related species of the *Oscheius* genus (enriched in A and B but depleted in D) was also significantly contributing to the taxonomic restructuring. However, most contrasting responses contributing to the plot-specific pattern of community restructuring were associated with the 17 genera that were significantly enriched or depleted in only a single transect relative to the reference transect E.

Despite this strong plot-specificity pattern, 9 genera showed consistent enrichment relative to transect E in at least two converted plots, forming a shared agroforestry signal. This signature includes tree taxa such as *Olea* (enriched in A, B, C, D) and *Fraxinus* (A, B, D), as well as herbaceous plants *Mercurialis* (B, C, D), *Triticum* (A, B, C) and *Medicago* (A, D). These plant enrichments were accompanied by increases in several associated taxa, including the arbuscular mycorrhizal fungi *Rhizophagus* (A, D) and the ecologically flexible fungi *Alternaria* (A, B, C). In addition, the ant *Lasius* (B, C) and the beetle *Brachypterus* (A, D) were consistently enriched, confirming patterns observed earlier in above-ground insect biodiversity monitoring. Conversely, five genera also showed consistent depletion relative to transect E in at least two converted plots: the plant genus *Parietaria* (B, D), and four fungal genera including the soil saprotroph *Aaosphaeria*, the metabolically flexible saprotroph *Talaromyces*, and the insect-associated fungi *Metarhizium*, all consistently depleted in converted plots (A, B, C, and D), as well as the wood saprophyte and putative plant pathogen *Clonostachys* fungi (depleted in A and D). Importantly, these 14 genera showing coherent shifts across multiple transects correspond to the most abundant taxa in the dataset and were all validated through direct field observations or ecological plausibility assessments (**Tables S3–S5**).

Together, these results indicate that agroforestry transition is mainly associated with plot-specific restructuring of multitrophic soil communities but with evidence of a restricted but consistent and abundant ecological signature, largely driven by vegetation diversification, specific enrichment of key symbiotic and saprotrophic fungi and increased insect diversity. Further monitoring will be required to confirm the stability of this emerging agroforestry complex signature over time.

### Soil functional diversification reveals plot-specific nutrient processing strategies across agroforestry transects

Given the very high bacterial richness observed along the agroforestry gradients of the farm but weak trends in their taxonomic restructuring, a functional protein annotation approach was explored to compare the bacterial functional potential of the soil and enrichment profiles across the agroforestry plots.

Metagenomic sequences were therefore assembled, generating 212,091 contigs across all soil samples (**Table S7**). From these assemblies, 257,751 protein-coding genes were predicted with a similar average length (496.4 ± 262.6 bp) and comparable length ranges across all soil samples (200–5,754 bp) suggesting similar assembly efficiency. Of these predicted proteins, 82,054 sequences could be taxonomically classified, of which 67,456 (82.2%) were assigned to bacterial proteins. To reduce redundancy among homologous sequences and minimize stochastic variation associated with individual protein predictions, proteins were further clustered into representative protein families. Using a minimum cluster size of three homologous proteins, 988 representative protein families were identified across the farm. Among these, 889 families (89.9%) were associated with at least one conserved protein domain based on Pfam annotation and to 55 functional categories derived from Gene Ontology (GO terms). A relatively constant fraction of uncharacterized protein families (10.0 ± 1.0%) was observed across all soil samples, indicating the presence of a small but consistent pool of potentially novel bacterial functions.

To characterize the functional potential of soil bacterial communities along the farm transects, an intersection analysis was performed using 253 unique Pfam–GO pairs linking protein structural domains to functional annotations (**Figure 6a**). This analysis revealed a restricted functional repertoire shared across plots, together with a highly heterogeneous distribution of transect-specific functions. Only 28 functional units were common to all five transects and mainly corresponded to the core molecular machinery required for bacterial gene expression, including ribosomal proteins (e.g. PF00164, PF00203, PF00238), RNA polymerase subunits (PF04997, PF04560), sigma transcription factors (PF04539, PF04542), and the elongation factor Tu GTP-binding domain (PF00009). Beyond this conserved core, transect-specific functional repertoires were observed: Transect A exhibited the largest number of unique functions (81 Pfam–GO units), including nutrient transport systems (sugar, ammonium, and phosphate transporters; PF00083, PF00909, PF01384), carbohydrate-processing enzymes such as α-amylase (PF00128), and several regulatory and stress-response proteins, including AraC and LuxR transcriptional regulators (PF00165, PF00196) and AhpC/TSA antioxidant proteins (PF00578), indicating a putative enhanced capacity for nutrient acquisition and environmental response. In contrast, transects B and C displayed smaller sets of specific functions, including the carbon starvation protein CstA (PF02554) and flavin oxidoreductases (PF01593) in B, and glycosyltransferases (PF00953) in C. No functional units were uniquely associated with transect D, suggesting a largely overlapping functional repertoire with the other plots, but also reflecting the smaller number of assembled protein families in this transect. Finally, the reference transect E showed only a few specific functions, mainly related to central metabolism and replication, including DNA polymerase III (PF07733) and the malic enzyme NAD-binding domain (PF03949). Together, these results indicate that soil bacterial communities share a conserved housekeeping functional backbone, while functional diversification occurs through localized enrichment of metabolic and regulatory functions, especially evident in the most advanced agroforestry plot (transect A).

**Figure 6.**
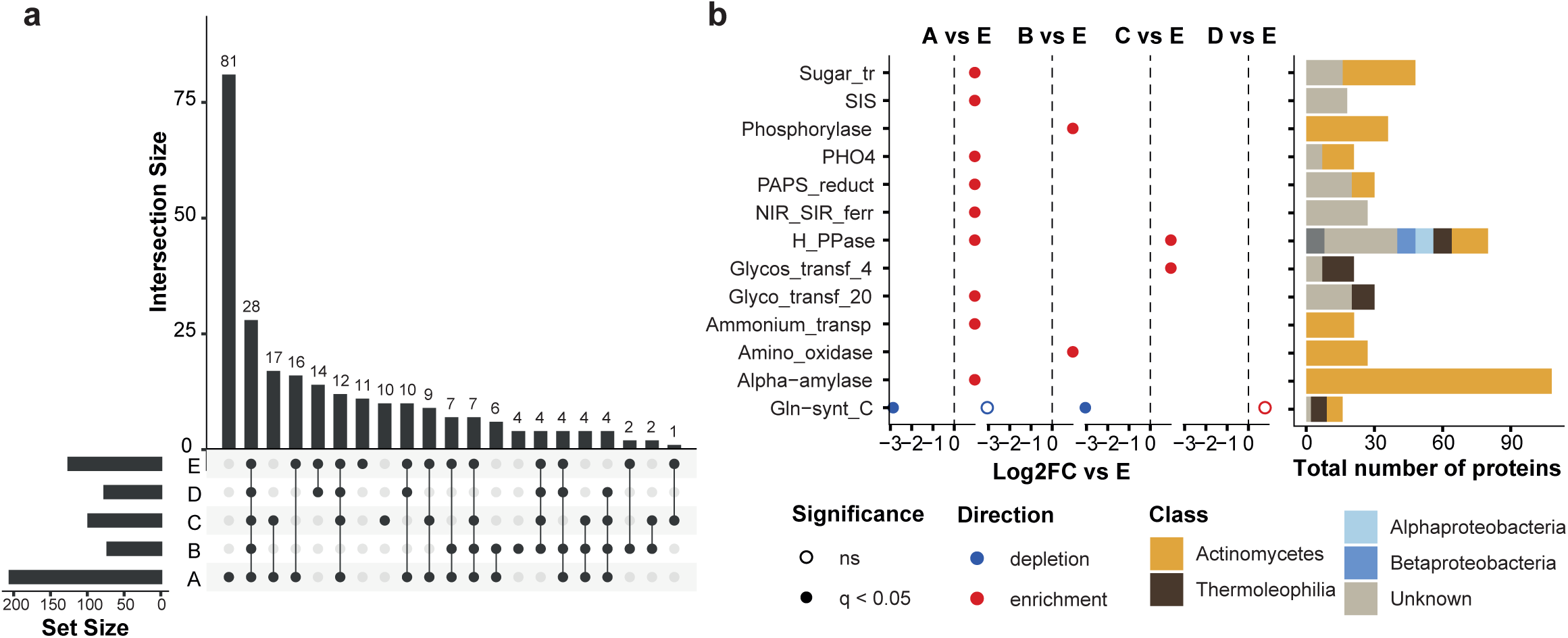
Functional overlap and nutrient-related functional shifts across agroforestry transects. **(a)** UpSet diagram showing the distribution and overlap of the 253 functional units defined by unique Pfam–GO associations across the five transects (A–E). Horizontal bars on the left indicate the number of functional units detected within each transect. Vertical bars represent the number of functional units shared among the transect combinations indicated by the connected points below each bar. **(b)** Left panel: Differential abundance of selected nutrient-related Pfam families across transects (A–D) relative to the reference plot (E). Each point represents a Pfam family, with its position on the x-axis indicating the log2 fold change (Log2FC) in abundance relative to transect E. Right panel: Cumulative counts of proteins associated with each nutrient-related Pfam family across all samples, colour-coded by taxonomic class.

To determine whether this functional diversification also translated into a significant enrichment of the soil nutrient-processing capacity, a targeted search for protein families associated with the acquisition and/or metabolism of carbohydrates, nitrogen, phosphorus, and sulfur was conducted (see Methods). This analysis validated 18 Pfam domains (**Table S8**), 13 of which showed significant enrichment in at least one transect (Fisher’s exact tests, FDR-adjusted q < 0.05), with the greatest enrichment observed in the most advanced agroforestry plot (**Figure 6b**). Indeed, transect A exhibited a coordinated enrichment of functions involved in carbohydrate turnover and transformation, notably through sugar transporters (PF00083), alpha-amylases (PF00128), the SIS domain (PF01380), and glycosyltransferase family 20 (PF00982). This pattern was accompanied by enrichment of ammonium transporters (PF00909), phosphate transporters (PF01384), and markers of sulfur assimilation (PAPS reductase, PF01507), suggesting an increased potential for inorganic nutrient mobilization. In contrast, the intermediate transects exhibited more limited responses. Transect B was characterized by significant enrichment of enzymes mobilizing intracellular carbon and nitrogen, such as carbohydrate phosphorylases (PF00343) and flavin-containing amine oxidoreductases (PF01593), while transect C showed enrichment of stabilization functions for carbon and phosphorus (glycosyltransferase family 4, PF00953, and H⁺-pyrophosphatase, PF03030), the latter representing the only protein domain shared with transect A. Transect D showed minimal differentiation from the unconverted plot, with the exception of glutamine synthetase (PF00120), which was enriched in D but significantly depleted in A and C, suggesting contrasting nitrogen assimilation strategies between plots rather than a uniform functional change. Finally, nutrient-related functions were primarily associated with protein families from Actinomycetes, consistent with their dominance within the soil bacterial community (**Figure 4a**), while specific functions such as glycosyltransferases (PF00953, PF00982) and H⁺-pyrophosphatase (PF03030) also involved bacteria from the Thermoleophilia and Proteobacteria classes.

Overall, these results confirm that agroforestry transition induces non-uniform, plot-specific changes in microbial functional potential. While transect A shows the most marked improvement in nutrient uptake and mobilization capacity, the other transects exhibit alternative and partially redundant functional strategies, consistent with the previously observed heterogeneous restructuring of the community.

## Discussion

This study provided an integrated assessment of above- and below-ground ecological responses to agroforestry transition within a Mediterranean perennial orchard. By combining adaptive field monitoring with soil shotgun metagenomics across a farm-scale conversion gradient, agroforestry was shown to be associated with increased insect richness and functional complexity above ground, spatial reorganization of insect-mediated services, and non-linear restructuring of soil communities below ground. Rather than following a uniform or gradual trajectory, the transition was characterized by functional diversification of soil communities, strong plot-specific responses, and the emergence of a restricted but consistent agroforestry signature across multiple trophic levels.

Above ground, the agroforestry transition was associated with a significant increase in insect richness and functional complexity. Across transects and months, insect richness increased significantly with flowering plant richness, supporting a resource-based mechanism whereby greater floral diversity expands feeding, nesting, and refuge opportunities for insects. This pattern is consistent with agroecological theory predicting that diversified vegetation enhances habitat heterogeneity and trophic interactions in agroecosystems (Altieri, 1999; Landis et al., 2000). In our system, the unconverted orchard largely contained a subset of the insect morphogroups recorded in converted plots, whereas the most mature agroforestry plots supported additional groups, including beetles, true bugs, aphids, crickets, and grasshoppers, alongside consistently present pollinators. This suggests community expansion rather than taxonomic replacement. This expansion of insect communities without apparent replacement of dominant pollinators is consistent with studies reporting that diversified agricultural systems can support additional arthropod guilds while maintaining pollination functions and increasing ecological resilience (Rader et al., 2015; Vialatte et al., 2025)

Importantly, the increase in insect diversity did not correspond to greater pest pressure on crops. Crop flowers were visited almost exclusively by beneficial pollinators, whereas potentially harmful taxa were mainly associated with supportive and spontaneous vegetation and rarely interacted with crop reproductive structures. This spatial segregation suggests that non-crop vegetation may buffer pest activity by providing alternative habitats or resources away from crops while maintaining pollination services. Similar patterns have been reported in diversification-based agricultural systems, where increased vegetation heterogeneity and temporal continuity of resources are associated with shifts in arthropod community composition, greater abundance of beneficial taxa, and reduced pest damage (Seimandi-Corda et al., 2026; Vialatte et al., 2025). From a farm management perspective, these patterns suggest that the timing and intensity of grass cutting could influence the ecological buffering capacity of spontaneous vegetation. Delaying mowing during crop flowering or until the seasonal emergence of predator communities may help maintain alternative habitats for both pollinators and natural enemies. Likewise, identifying which spontaneous or supportive plant species preferentially attract potentially harmful taxa could inform targeted planting strategies aimed at spatially diverting pest activity away from crops. Although these interpretations remain exploratory, they illustrate how adaptive biodiversity monitoring could support more ecologically informed farm management decisions.

In contrast to the pronounced above-ground patterns, soil communities remained taxonomically very similar across the farm. At the farm level, bacterial communities were strongly dominated by Actinomycetota, a pattern commonly reported in Mediterranean soils, where their distribution appears closely linked to geomorphometric conditions (Pelacani et al., 2025). Most genera were shared among all plots, and overall richness changed little along the agroforestry transition. However, this apparent stability masked important shifts in the relative abundance of taxa; Rather than replacing existing soil organisms with entirely new communities, agroforestry transition in the farm mainly reshaped the relative abundance of already-present taxa. This pattern is consistent with ecological studies showing that land-use change, under neutral soil pH, often alters microbial community structure through stochastic and local processes rather than through major species turnover (Goss-Souza et al., 2017; Tripathi et al., 2018). Similarly, as shown by (Sveen et al., 2025), showed that ecosystem development altered the functional diversity and nutrient-cycling capacities of microbial communities more strongly than their taxonomic composition during grassland-to-forest succession. In our study, these shifts likely reflected differences in vegetation composition, litter accumulation, root activity, and local management history, which progressively modified resource availability and microhabitats within each plot. Above-ground diversification may also have indirectly influenced microbial functional capacities through changes in organic matter inputs and substrate heterogeneity. Consistent with this interpretation, other studies identified litter quality and soil pH as major drivers of microbial community structure and functioning during ecosystem succession (Sveen et al., 2025). Consistently, soil communities in our study did not evolve along a single linear trajectory with time since conversion, but instead followed plot-specific paths, suggesting that agroforestry systems can develop through multiple ecological trajectories depending on local conditions, litter quality and management practices (Louisson et al., 2024)

Despite the strong plot-specific restructuring of soil communities, a restricted but coherent agroforestry signature emerged across several converted plots. This shared signal included the enrichment of herbaceous genera such as *Mercurialis*, *Medicago*, and *Triticum*, together with the depletion of *Parietaria*, indicating a shift toward more diversified and managed ground vegetation. Below ground, this vegetation signal was associated with enrichment of the arbuscular mycorrhizal fungi *Rhizophagus*, a genus commonly associated with enhanced nutrient uptake and plant productivity in diversified agroecosystems (Mwampashi et al., 2024; Wu et al., 2023). In contrast, the enrichment of the fungi *Alternaria*, particularly *Alternaria atra*, was more ambiguous. Although several Alternaria species are recognized plant pathogens, recent work on *Alternaria atra* suggests substantial lifestyle plasticity, with isolates functioning as endophytes, saprotrophs, or pathogens depending on ecological context (Schmey et al., 2026; Woudenberg et al., 2015). The limited disease symptoms observed in the field, together with independent leaf analyses showing only trace detection of *Colletotrichum* and no evidence of *Alternaria* infection (**Table S3**), suggest that the *Alternaria* DNA found in soil may primarily reflect residual soil DNA or a non-pathogenic ecological role within the system. Interestingly, the diversification of insect communities observed above ground was also reflected in soil-associated taxa, with enrichment of *Lasius* ants and *Brachypterus* beetles across converted plots, suggesting stronger links between vegetation, litter, and soil habitats. Conversely, agroforestry plots consistently showed depletion of the fungi *Metarhizium*, *Talaromyces*, *Clonostachys*, and *Aaosphaeria*, taxa associated with diverse ecological strategies including saprotrophy, insect association, and opportunistic colonization(Frisvad et al., 2013; Zimmermann, 2007). The decline of *Metarhizium*, an insect-associated entomopathogenic fungus, may reflect reduced dependence on soil-borne pest regulation following the spatial redistribution of pest activity and the emergence of more diverse predator communities (Behie et al., 2012). Together, these patterns suggest that vegetation diversification progressively reorganized trophic interactions and nutrient-processing pathways across above- and below-ground compartments, leading to the emergence of an integrated multitrophic agroforestry signature.

Functional analyses of soil bacterial communities further showed that agroforestry effects were more visible at the level of metabolic potential than taxonomic diversity. Although bacterial richness and composition changed only modestly, predicted protein functions varied substantially among plots. The most mature agroforestry plot showed coordinated enrichment of functions related to carbohydrate turnover, nitrogen and phosphorus transport, sulfur assimilation, and environmental response, suggesting greater capacity for nutrient acquisition and transformation. In the converted plots, organic matter inputs originated largely from in-field processes, including pruning residues, chop-and-drop management, wood-chip deposition, and cut herbaceous vegetation. These diversified biomass inputs likely increased substrate heterogeneity and carbon availability, potentially favoring saprotrophic activity, reshaping mycorrhizal associations through greater root diversity, and contributing to the enrichment of nutrient- and carbon-cycling functions observed in converted plots. This interpretation is consistent with studies showing that plant-derived inputs such as root exudates and litter, together with cropping practices, can drive distinct microbial metabolic responses and nutrient-cycling strategies particularly regarding C-N-P-cycling genes and metabolic specialization across agricultural systems (Krause et al., 2025; Seitz et al., 2024; Sveen et al., 2025; Yan et al., 2018). Intermediate plots showed more restricted or specialized functional responses, whereas the most recent conversion remained closer to the reference plot, indicating that functional changes emerged unevenly across the transition. The observed decoupling between relatively stable taxonomic structure and variable functional potential is consistent with the concept of microbial functional redundancy, whereby different microbial assemblages can maintain overlapping ecological functions while reorganizing dominant metabolic pathways in response to local resource conditions (Louca et al., 2018). Together, these patterns suggest that agroforestry transition altered nutrient-cycling strategies rather than major bacterial community replacement. The decoupling between taxonomic stability and functional diversification points to substantial functional plasticity within the soil microbiome.

Taken together, these findings highlight the importance of vegetation diversification as a practical management lever linking above- and below-ground ecological processes. From a farmer perspective, maintaining diverse spontaneous and supportive vegetation may not only sustain pollinator communities but also spatially buffer pest activity and promote more functionally diverse soil systems. The strong plot-specific responses observed across the farm further suggest that agroforestry transition does not follow a single optimal trajectory, but instead depends on local management history, planting strategies, and ecological context. This reinforces the importance of adaptive and farmer-centred monitoring approaches focused on functional trends and early warning signals rather than fixed biodiversity targets. In this regard, the present study also illustrates the value of close collaboration between farmers, agronomists, and scientists, as the experimental design itself emerged from an existing long-term agroforestry transition initiated and managed at the farm level. Such co-constructed approaches are increasingly recognized as essential for developing realistic and context-adapted agroecological research and innovation frameworks (Duru et al., 2015; Tittonell, 2014).

Several limitations should nevertheless be considered when interpreting these results. The study was conducted on a single farm and relied on a chronosequence design, limiting broader generalization and preventing full separation of temporal effects from plot-specific histories. Insect abundance estimates remained semi-quantitative, and monitoring was restricted to spring, potentially overlooking important seasonal dynamics outside the main flowering period. In addition, pest observations were mainly restricted to visible interactions with flowering structures, potentially underestimating low-density or canopy-associated infestations within crop trees. Similarly, metagenomic functional profiles reflect potential rather than realized microbial activity and therefore cannot directly infer active ecosystem processes, while the limited spatial and temporal soil sampling may have underestimated within-plot heterogeneity. Future work should extend this approach across larger spatial scales, multiple farms, and longer temporal periods while integrating direct measurements of nutrient fluxes, decomposition, and trophic interactions. Replicating this type of integrated monitoring framework across contrasted agroforestry systems will be essential to determine whether the multitrophic signatures observed here represent generalizable features of agroforestry transition or context-specific trajectories.

Despite these limitations, this study provides empirical evidence that agroforestry transition can enhance functional biodiversity and reorganize ecological interactions at the farm scale without necessarily increasing overall soil taxonomic richness. By integrating above- and below-ground perspectives, it highlights how vegetation diversification can simultaneously influence insect dynamics, pest spatial distribution, and soil functional potential. The emergence of functionally diverse insect communities, the apparent buffering of pest activity by non-crop vegetation, and the localized enhancement of nutrient-processing functions together suggest that agroforestry can support the development of more resilient and self-regulating production systems. More broadly, our findings align with emerging ecological perspectives emphasizing the importance of understanding biodiversity, ecosystem functioning, and species interactions as interconnected components of sustainable agricultural transitions (Windsor et al., 2022). Such integrative approaches may become increasingly important for guiding the design and monitoring of future regenerative farming systems.

## Methods

### Study site and farm management

The Southern Lights Farm (36°50′37.7″ N, 22°37′32.2″ E) is located in the southern Peloponnese region of Laconia, Greece. The farm lies within the “*Mediterranean South*” environmental zone as defined by (Metzger et al., 2005), with a mean annual temperature of approximately 19 °C and mean annual precipitation of 700–800 mm (HNMS, n.d.). The farm is located within an agricultural zone surrounded by conventionally managed olive and citrus monocultures (**Supplementary Fig. S1**). The broader landscape is characterised by relatively humid lowland conditions, including swamps and shallow groundwater. Soils are formed from calcareous parent material with marine influence, and the dominant soil texture on the farm is sandy clay.

Southern Lights Farm covers 3.2 ha and has been managed organically since its establishment in 1980, when the original olive trees were interplanted with citrus trees. The main crops currently include olives (Koroneiki variety), lemons, limes, mandarins and oranges (Navel, Merlin and Valencia variety). Since 2015, the farm has been irrigated every 7 days from late April to late September (depending on the weather conditions). Ground vegetation is mechanically cut approximately three times per year. Olive trees are pruned annually after harvest (November–December), while citrus trees are pruned every second year. Since the establishment of the farm, only manure was added yearly during the first years and then every 3 to 4 years. No other external soil inputs (fertilisers or amendments) have been applied.

### Agroforestry transition and planting system

A progressive transition from organic olive and citrus production to syntropic agroforestry began in 2015, initially motivated by the development of a multifunctional food forest system. This conversion aimed to establish a dense, highly diversified agroforestry system based on low input and high-biomass management approach to enhance soil quality, microclimate regulation, and long-term autonomy with minimal external dependencies and labour requirements. As different areas were converted at different times and with different degrees of diversity, the farm currently represents a gradient of agroforestry maturity. Five plots of comparable size (0.4 ± 0.2 ha) were delineated based on the starting year of conversion: plots A and B (converted in 2015), plot C (2018), plot D (2024), and plot E (not yet converted) (**Figure 1A**).

In plots A and B, most of the existing citrus rootstocks were re-grafted in 2015 to increase crop diversity and provide additional marketable citrus varieties, while fruit trees (mulberry, fig) naturally emerging from sudden sunlight exposition, were retained. Between 2016 and 2017, approximately 350 fruit trees representing 30 species, including tropical and non-native species, together with 200 support species were inter-planted. More support species were supplemented to the ones that didn’t survive in the following years. A multispecies cover crop mixture was introduced in 2019 and subsequently maintained through self-seeding and spontaneous species. In plot A, additional citrus re-grafting was conducted in 2023. In plot C, seven citrus species were grafted onto existing rootstocks in 2018, accompanied by additional supportive tree plantings. Plot D represents the most recent and most systematically implemented agroforestry conversion, with supportive tree planting and ground vegetation management introduced in 2024 to promote early-stage biomass production and ecosystem development. Plot E remains under organic orchard management without formal agroforestry conversion, serving as a non-converted reference within the farm. In the converted plots, organic matter input relies on in-field processes, including pruning residues, chop-and-drop management, surface deposition of wood chips, and cut herbaceous vegetation.

Across all plots, agroforestry was implemented as a linear system retrofitted onto the original farm layout. Trees were arranged in rows spaced 5 m apart, with main crop trees planted at 4 m intervals within rows. Support species were interplanted between the crop trees at approximately 30 cm distance, creating close spatial associations intended to enhance structural complexity, microclimate buffering, and in-field biomass production. The farm’s flat topography ensures relatively uniform surface water movement and comparable hydrological conditions across plots.

### Transect-based monitoring

An adaptive transect-based protocol was implemented to capture variation in floral resources and associated insect activity within the farm, with the aim of providing a comprehensive and management-relevant overview of plant–insect dynamics. The protocol prioritised exhaustive monitoring of all observable species along fixed transects rather than strict standardisation of survey duration, with the objective of detecting seasonal patterns and potential risk windows. Therefore, observation duration varied with floral resource availability, while spatial replication through repeated surveys of the same transects constituted the primary standardisation criterion.

Monitoring was conducted once per month from March to June 2024, under sunny conditions between 09:00 and 13:00. A fixed transect was established in each plot, with transect length adapted to plot size and a constant width of 2m. Each transect was subdivided into 3 m sections, and the time required to complete each transect walk was recorded. Abiotic conditions, including ambient temperature and wind intensity were also recorded. Each month, two consecutive transect surveys were conducted: (i) a plant community survey recording all visible plant and insect taxa along the transect, and (ii) an insect visitation survey specifically documenting insects and insect interactions with flowering plants. Flowering plant–insect interactions were defined as any direct contact between an insect and floral or immediately adjacent reproductive structures, including the occupation of floral tissues or stems by insects, with potential effects on plant viability and fruiting. Insects observed on vegetation or within transect sections that did not meet these criteria were nevertheless recorded and classified as outlier interactions. For each 3 m section, the frequency of an interaction or observation was recorded as the number of insect individuals per plant species; counts up to ten individuals were recorded directly, while higher abundances were conservatively capped at ten, resulting in semi-quantitative abundance estimates. This approach was adopted to maintain consistency with the farmer-oriented objective of capturing relative changes in insect activity across seasons rather than exact population sizes. Both sides of the transect were observed concurrently as observers progressed, to avoid double counting.

Finally, soil surface pH was measured with *Kraft & Dele Professional KD11408* portable soil detector at the beginning and end of each transect, while a 3 point (beginning, middle, and end) composite soil sample was collected at 20 cm depth along each transect in March for subsequent soil biodiversity analyses. To prevent cross-contamination, all sampling equipment was cleaned with 70% ethanol between samples.

### Species identification and classification

Both plant and insect species were identified based on morphological characteristics. When field identification was uncertain, species determination was validated using photos and web-based identification tools (www.inaturalist.org).

Each plant species was classified according to three complementary descriptors. First, species were categorised according to their functional role within the farm management system: crop species, corresponding to the primary cultivated trees managed for production; support species, including plants intentionally introduced or encouraged to support ecosystem functions such as biomass production, soil cover, microclimate regulation, or secondary yields; and spontaneous species, defined as plants establishing without direct human introduction and not systematically removed. Second, plant species were assigned to one of five agroforestry vertical strata based on their typical mature stature and spatial position within the system: emergent, corresponding to the tallest trees forming the upper canopy; high, including dominant canopy trees; medium, comprising sub-canopy trees and large shrubs occupying intermediate vertical space; low, including shrubs and small woody species; and ground cover, encompassing herbaceous species, which included both low-growing and some climbing taxa. Finally, all species were categorised by growth form as tree, bush, or herbaceous, independently of their functional role and vertical stratum.

Insect taxa were identified to species or genus when possible or kept grouped into operational morphogroups. Insects were also assigned to relevant ecological role categories: beneficial pollinators included insects that contribute to the pollination of flowering plants, thereby supporting fruit or seed production; natural predators comprised insects that prey on herbivorous taxa, contributing to pest regulation and potential pests included insects known or suspected to negatively affect crops through feeding on vegetative or reproductive plant tissues. Species whose roles varied across life stages or contexts (e.g. pollinating adults with predatory or parasitic larvae, or ants exhibiting both pest facilitation and predation) were assigned to the mixed category. Finally, insects with no documented or negligible direct influence on crop pollination, pest regulation, or damage were classified as having a limited role in crops.

### Insects diversity analyses and statistics

Diversity indices were calculated at the transect × month level using the *vegan* package in R. Species richness was computed from presence–absence data derived from species-level identifications for both flowering plants and insects, where a species was considered present if observed at least once within a transect and month. To assess changes in insect community structure, Shannon diversity indices were calculated using frequency-weighted abundance data, where species abundances were derived from the semi-quantitative estimate of flowering plant–insect monitoring.

Association between flowering plant and insect species richness was tested using linear mixed-effects models fitted with the lmer function in the *lme4* package in R. Month was included as a fixed effect to account for seasonality, and transect identity was included as a random intercept to account for repeated sampling within transects. Insect compositional differences were analysed at the morphogroup level using presence–absence data using Jaccard dissimilarity. Chord diagrams were generated using the R package *circlize* to visualise frequency-weighted plant–insect interaction networks. Analyses focused on descriptive characterisation of interaction patterns; formal statistical testing of network structure was not conducted at this stage.

### Soil sampling and DNA extraction

A total of five composite soil samples were collected, one along each transect during the first monthly assessment in March 2024 and stored at −20 °C until DNA extraction. DNA was isolated from 250 mg of each soil sample, strictly following the manufacturer’s protocol provided with the QIAGEN DNeasy PowerSoil® kit. DNA was eluted from the column with a 25-minute incubation. During DNA extraction, ZymoBIOMICS Microbial Community Standards (D6310; log distribution) were included to assess the qualitative and quantitative performance of the extraction and to serve as an internal control for sequencing. Extracted DNA was stored at −20 °C until library preparation.

### Library preparation and DNB-based sequencing (MGI)

A total of six DNA libraries (5 soil samples and one community standard) were prepared using the MGIEasy Fast FS Library Prep Set (version 2.0; MGI Tech Co., Ltd.) following the Fast FS Library Prep Set User Guide (version 1.0). Depending on the available input DNA, either 13.5 ng or 230–270 ng were used, with fragmentation times set to 18 min and 12 min, respectively, followed by single-size selection. Prior to adapter ligation, UDB adapter dilution and endpoint PCR cycle numbers were determined according to the initial DNA input, as specified in the manufacturer’s guidelines. Specifically, six PCR cycles were applied to libraries prepared from 230–270 ng of input DNA, whereas 10 PCR cycles were applied to the library prepared from 13.5 ng of input DNA.

Library quality was assessed using the Agilent TapeStation 4150 with the High Sensitivity D1000 assay kit, and library concentration was measured using the Invitrogen Qubit 4 Fluorometer with the Qubit dsDNA HS Assay. Equimolar amounts of libraries were pooled to a total of 1 pmol. Pooled libraries then underwent DNA denaturation, single-stranded circularization, enzymatic digestion, digestion product cleanup, and quality control steps according to the Fast FS Library Prep Set protocol. Subsequently, 45 fmol of single-stranded circular DNA were converted into DNA nanoballs (DNBs) by rolling circle amplification. The resulting DNBs were loaded onto 4 different flowcell lanes (technical replicates) and sequenced on a DNBSEQ-G400 platform at the Genomics Facility of BSRC “Alexander Fleming” using the G400 FCL PE150 High-throughput Sequencing Set (MGI Tech Co., Ltd.), following the DNBSEQ-G400RS High-throughput (Rapid) Sequencing Set User Manual (version 8.0).

### Sequence quality control and trimming

QC analysis of the resulting 24 libraries was performed using FastQC (version 0.12.0) and included assessments of per base quality, per sequence quality scores, per base sequence content, per base GC content, per base N content, sequence length distribution, sequence duplication levels, overrepresented sequences, and k-mer content (Andrews S., 2010). Subsequently, libraries were trimmed using bbduk tool (version 39.80) (Bushnell, 2014). This included trimming specific MGI adapters and JGI library contaminants, which contains sequences from common contaminants such as vectors, adapters, primers and human sequences. Different k-mer values (k=5, 8, 10, 12, 15, 20, 30) were tested. Ultimately, default parameter k=27 was used, which provided the best balance between trimming efficiency and quality retention. After trimming, FastQC was rerun to ensure the samples met high-quality metrics, confirming that the chosen k-mer parameter effectively removed contaminants while maintaining sequence integrity. Paired reads were interleaved in a single file to ensure that taxonomy classification forward and reverse reads were classified together.

### Taxonomic classification and sequencing depth assessment

Taxonomic classification of all libraries was performed using Kraken2 (version 2.1.3) against the NCBI nucleotide (nt) database with a confidence threshold of 0.05 (Wood & Salzberg, 2014). This threshold significantly assessed both qualitative and quantitative bacterial composition of the libraries derived from ZymoBIOMICS Microbial Community Standards (**Supplementary Fig. S3a**). Because nt-NCBI taxonomy frequently includes provisional names (e.g. “*Candidatus*” or “candidate”), taxonomic annotations were standardized by flagging provisional names across taxonomic ranks and reconstructing Kingdom-level assignments from phylum-level taxonomy. Species-level abundance tables were generated by aggregating read counts per species across the 20 soil libraries, resulting in a species-level OTU table comprising 18,075 taxa. Sequencing depth and library comparability were assessed by constructing species-level rarefaction curves using the rarecurve() function from the *vegan* R package, with rarefaction performed at the minimum sequencing depth across libraries. This minimum sequencing depth showed that species accumulation curves approached a plateau for most libraries, indicating that species richness estimates were overall stable across the libraries (**Supplementary Fig. S3b-c**).

### Metagenomic data and statistical analyses

All analyses were performed in R (v4.5.1). A phyloseq object was constructed from the OTU table, taxonomy table and sample metadata using the *phyloseq* package (McMurdie & Holmes, 2013). Taxa were agglomerated to the genus level using the tax_glom function. To reduce noise from extremely low-abundance taxa, genera were retained only if they reached a relative abundance of at least 0.01% in at least 15% of sequenced libraries across all dataset. After filtering, 1046 genera were retained for downstream analyses.

To assess soil taxonomic composition, Richness was calculated separately for major taxonomic groups (Bacteria, Archaea, Fungi, Plantae, and Invertebrates) across transects. Invertebrates were defined by grouping genera belonging to the phyla Arthropoda, Nematoda, and Mollusca. Genus richness was calculated using the specnumber() function from *vegan* package and summarised at the transect level as mean ± standard error across technical replicates. Genus prevalence was assessed and quantified using presence–absence–based Jaccard dissimilarity, calculated with the vegdist() function from *vegan*. Linear models describing the association between genus richness and years since agroforestry conversion were fitted.

To assess soil taxonomic dynamics, Shannon and Simpson diversity indices were calculated on relative abundance data using the diversity() function from *vegan*. Pielou’s evenness was calculated as Shannon diversity divided by the natural logarithm of genus richness. All indices were summarized at the transect level as mean ± standard error across technical replicates. Beta diversity was quantified using Bray–Curtis dissimilarity from relative abundance at the genus level. Global compositional structure was visualized using Principal Coordinates Analysis (PCoA) performed on the Bray–Curtis distance matrix using multidimensional scaling (cmdscale, k = 2). To quantify pairwise compositional differences, mean Bray–Curtis dissimilarity was calculated for all transect pairs by averaging pairwise distances among all corresponding libraries. Within-transect dissimilarities were used to estimate technical baseline variability.

Differential abundance analysis was performed at the genus level on taxa belonging to the kingdoms Fungi, Plantae, and Animalia (restricted to the invertebrate phyla Arthropoda, Nematoda, and Mollusca) using *ANCOMBC* package (Lin & Peddada, 2020) implemented on the phyloseq object. ANCOM-BC2 fits a bias-corrected log-linear model that accounts for differences in library size and compositional effects, producing bias-corrected log fold changes (lfc), standard errors, and Benjamini–Hochberg–adjusted q-values. Transect was included as a fixed effect, with the unconverted transect (E) set as the reference level. Structural zero detection (struc_zero = TRUE) and negative lower bound correction (neg_lb = TRUE) were enabled to improve stability of the estimates when taxa were absent in the reference group. Genera were considered robustly differentially abundant when they passed the ANCOM-BC2 sensitivity screening (passed_ss = 1), met a false discovery rate threshold (q < 0.05), and exhibited at least a two-fold change in abundance relative to the reference transect (|log2FC| ≥ 1). Differential abundance patterns were visualized as a heatmap of log2 fold changes using the *pheatmap* package in R, with hierarchical clustering applied to both genera and transects based on Euclidean distance.

### Field verification of metagenomic taxa

Metagenomic plant and invertebrate genera were validated and categorized using observations obtained during transect monitoring, management records and field observations reported by the farmer, and regional occurrence data for Greece and the Mediterranean region (e.g., iNaturalist and local biodiversity databases). Genera were classified into four verification categories: direct field verification, when observed visually during surveys or confirmed by the farmer; regionally plausible, when not observed in the field but known to occur naturally in the region; crops or supporting vegetation, when their presence is consistent with farm management practices or cultivated plant species; or unlikely/false positive, when the identification is ecologically improbable for the region or likely results from taxonomic misclassification in the bioinformatic pipeline. Except for false positive genera, the most abundant predicted species from the metagenomic dataset was extracted for each validated genus and, where possible, further verified through additional field observations or regional occurrence records. For fungal taxa with plant pathogenic potential, additional verification was attempted through field inspection of foliar disease symptoms and, when possible, laboratory analyses to confirm pathogen presence.

### Metagenomic bacterial protein annotation and functional enrichment analysis

Metagenomic reads underwent normalization via *BBNorm* (version 39.80) to mitigate data redundancy (Bushnell, 2014). The resulting reads were assembled into contigs using *MEGAHIT* (v1.2.9) (Li et al., 2016). By reconstructing reads into contigs, we established a more robust framework for predicting protein-coding sequences. This transition is essential, as the increased length of contigs facilitates more reliable Open Reading Frames (ORF) detection than is possible with individual reads. ORFs were predicted from assembled contigs using *prodigal* (version 2.6.3) (Hyatt et al., 2010) to identify putative protein-coding genes. Contigs were subsequently taxonomically classified using Kraken2 (Wood & Salzberg, 2014), and only contigs assigned to Bacteria were retained for downstream functional analyses.

Predicted protein from bacterial contigs were extracted and clustered into representative protein families using MMseqs2 (Steinegger & Söding, 2017) to reduce redundancy among homologous sequences. Only family clusters containing at least three proteins were retained. The representative sequence of each cluster was functionally annotated against the Pfam-A (version 37) database (Paysan-Lafosse et al., 2025) of profile Hidden Markov Models (HMMs), which contains 21,979 families and 709 clans . Pfam hits were identified using the hmmsearch tool from the HMMER 3.3.2 package31(Frith et al., 2010), applying the default trusted cutoff (Mistry et al., 2021). Proteins lacking PFAM annotations were classified as novel. Gene Ontology (GO) terms associated with PFAM domains were retrieved using InterPro2GO mapping from InterPro to define functional units corresponding to unique PFAM–GO associations. The distribution of functional units across transects was summarized using intersection analysis and visualized with *UpSetR* package (Conway et al., 2017; Lex et al., 2014).

Targeted analysis of nutrient-related bacterial functions was performed by screening protein families based on Pfam annotations and associated Gene Ontology (GO) terms using a keyword-based approach. Keywords were selected to capture functions involved in the microbial processing of key plant-relevant nutrients, including “Carbohydrate” (e.g. sugar transporters and glycosyl hydrolases), “Nitrogen” (e.g. ammonium transport and amino acid turnover),”Phosphorus” (e.g. phosphate transport and pyrophosphatases), and “Sulfur” (e.g. sulfate reduction pathways) categories. Resulting candidates were manually inspected to remove generic function and false positives. Enrichment of these curated Pfam families across transects was assessed using Fisher’s exact tests, with p-values adjusted for multiple comparisons using the Benjamini–Hochberg false discovery rate correction. Functional direction was determined based on log2 fold change.

## Data and code availability

The metagenomic data sets have been deposited in the Sequence Read Archive (SRA) from NCBI under BioProject ID PRJNA1470656. Biodiversity data, source codes and tables to reproduce all analyses and figures in this paper are available at Zenodo XXXX.

## Acknowledgements

We thank MGI Tech Co. Ltd for reagents and assistance; Bernard Okere of MGI Tech Co. Ltd and Vagelis Harokopos of [VARELAS S.A.](https://www.varelas.gr/) for support and guidance. We also thank Ashley Franks and Jennifer Wood from [La Trobe University](https://www.latrobe.edu.au/research/centres) for their initial guidance and methodological expertise in soil biodiversity analysis and reading the final version of the manuscript. We are grateful to [Regenerative Farming Greece](https://regenerativefarminggreece.org/), particularly Giuseppe Sannicandro, for training and discussions on agroforestry and soil ecosystems, and to Panos Darmos for sharing his knowledge and long-term vision for the Southern Lights Farm. We acknolwdge the bioinformatic support provided by Fotis Baltoumas and Alexandros Galaras. This work was supported by the [Hellenic Foundation for Research and Innovation](https://www.elidek.gr/en/) (H.F.R.I.) through the “3rd H.F.R.I. Research Grant to Support Postdoctoral Researchers” (7832-YGIAMEL), awarded to S.P., the “3rd H.F.R.I. Research Grant to Support Faculty Members and Researchers” (23592-EMISSION), and a research grant from the “Foundation Santé” awarded to G.A.P. The [Fleming Genomics Facility](https://www.fleming.gr/services/facilities/genomics) is supported by project SingleOut (HFRI-FM17C3-3780) funded by the Hellenic Foundation for Research and Innovation (H.F.R.I.), project MIS 6004752 funded by the Regional Operational Programme ’ATTICA’ (NSRF 2021–2027) and co-financed by Greece and the European Union (European Regional Development Fund), and the National Recovery and Resilience Plan Greece 2.0, funded by the European Union – NextGenerationEU.

## Author contributions

S.P. and S.D. designed the research. S.P., Ev.K., J.M. and P.V. acquired field data. S.P., P.L. and El.K. conducted wet laboratory experiments. S.P. and E.A. developed codes, analysed the data and provided data visualization. Ev.K., J.M. and M.C. provided field data validation. S.P. wrote the original draft. J.M., Ev.K., P.V., P.H. and S.D. reviewed and edited the manuscript. S.P. P.H. G.A.P and S.D. supervised the study and provided funds.

## Competing interests

The authors declared no competitive interests.

## Notes

### Competing Interest Statement

The authors have declared no competing interest.

