## Supplementary Materials for "Agroforestry transition increases insect diversity and reorganizes soil function in a Mediterranean orchard"

### Supplementary Figures

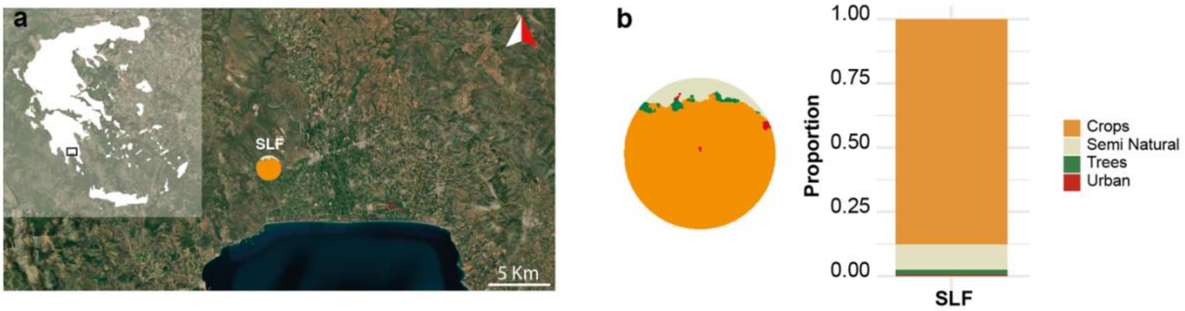


**Supplementary Fig. S1. Landscape context of the farm. (a)** Geographic location of the Southern Lights Farm (SLF), Greece within the surrounding landscape. **(b)** Landscape composition within a 1 km radius around the farm. The circular map illustrates land-cover classes derived from the 2018 CORINE Land Cover dataset, and the bar plot shows the proportional contribution of each class. Land cover was classified into four categories: crops (including horticulture, olive groves, and fruit crops), semi-natural grasslands (including abandoned agricultural fields, urbanized grasslands, and grasslands used for livestock), tree-dominated semi-natural vegetation (forests, hedgerows, shrubs), and urban areas. Land-cover classification and spatial analyses were conducted in QGIS, and landscape composition was quantified based on class proportions within the 1 km buffer around the farm.

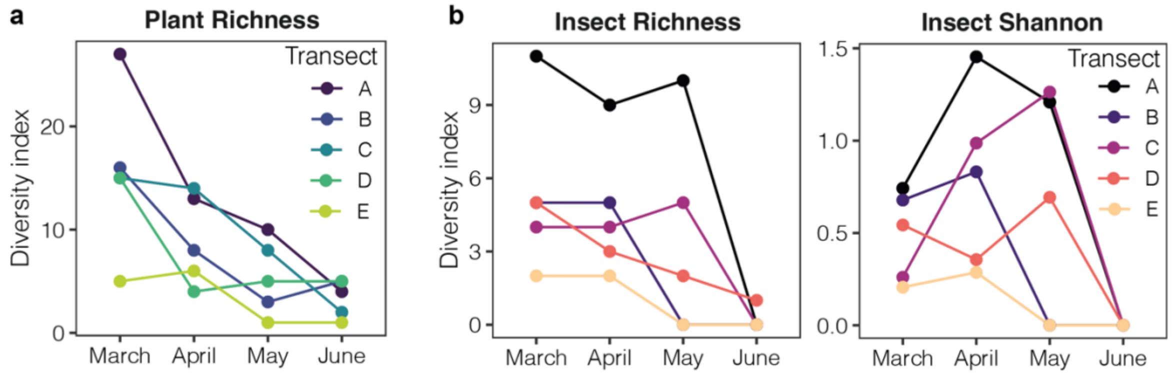


**Supplementary Fig. S2. Seasonal dynamic of flowering plant and insect diversity.** Monthly time plots showing the dynamic of plant (a) and insect (b) diversity indexes across transects. Richness was calculated using presence–absence data for insect and plant species, while insect diversity was quantified using a frequency-weighted Shannon index. Lines connect monthly values within each transect.

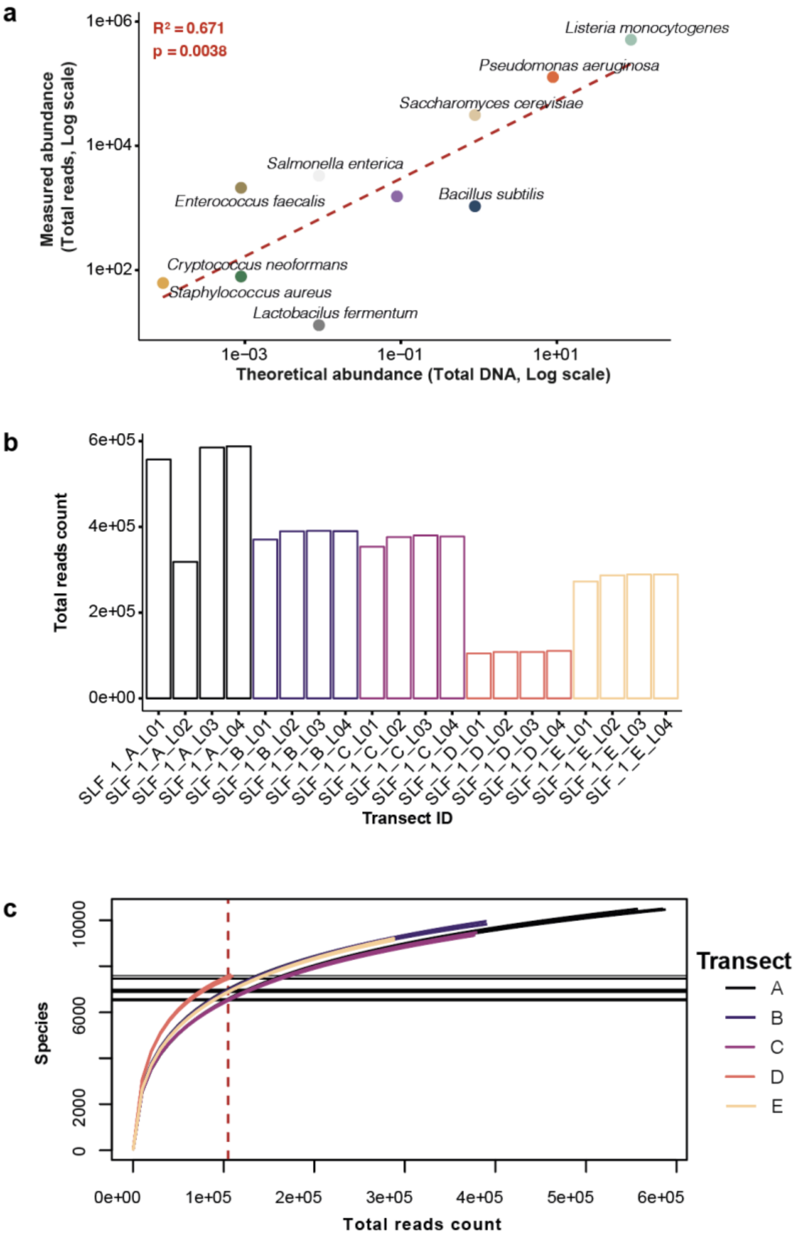


**Supplementary Fig. S3. Taxonomic classification and sequencing depth assessment. (a)** Theoretical versus measured abundance of the ZymoBIOMICS microbial community on a $\log_{10}$ scale. Each point represents a single species, with measured abundance expressed as total read counts and theoretical abundance derived from the manufacturer-provided community composition. The red dashed line represents a linear regression fitted on $\log_{10}$-transformed values with associated $R^2$ and p-value. **(b)** Barplots showing read count distribution across the sequenced libraries. **(c)** Species-level rarefaction curves showing species accumulation as a function of sequencing depth. Each curve represents one library. The vertical red dashed line indicates the minimum library depth (104,902 reads) used for standardization. Horizontal bands reflect the onset of a plateau in species discovery, where additional sequencing yields few new species.

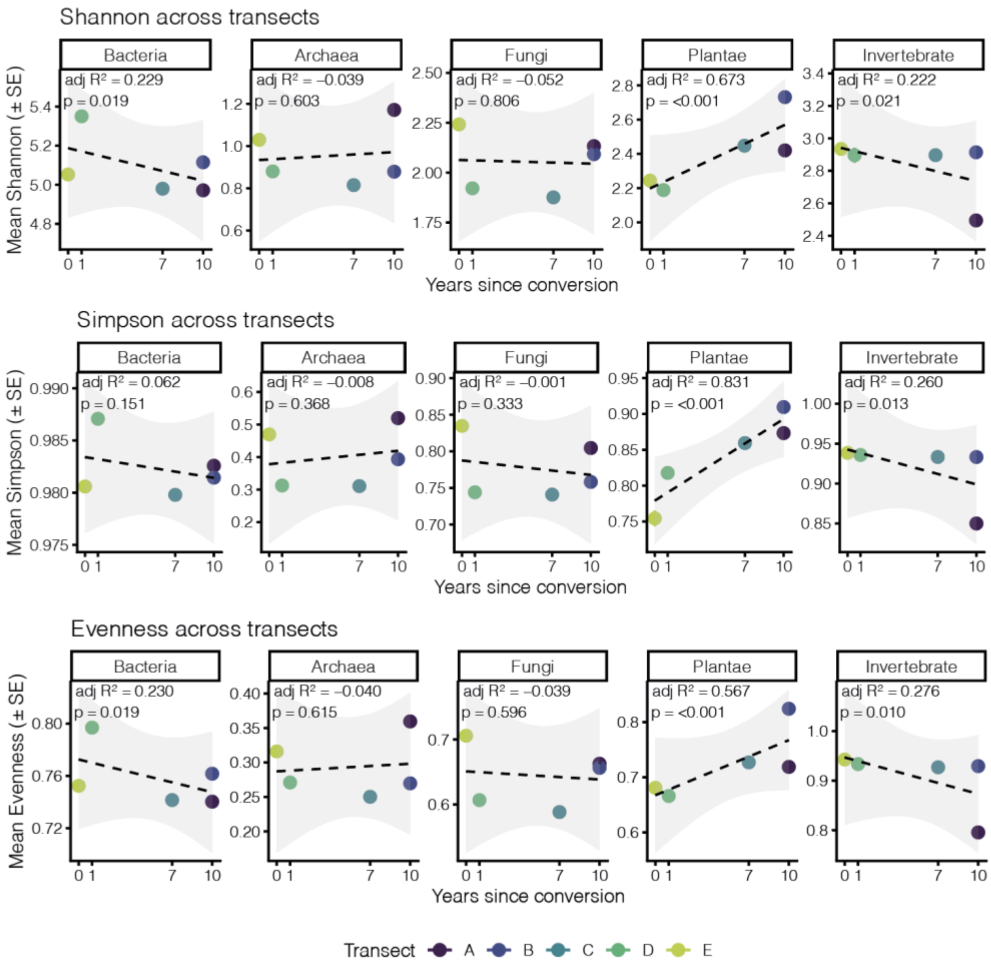


**Supplementary Fig 4. Temporal dynamics of alpha diversity across the farm.** Scatter plots showing Shannon diversity (top row), Simpson diversity (middle row), and Pielou's evenness (bottom row) for major taxonomic groups in relation to years since agroforestry conversion. Points represent mean diversity values ± standard error (SE) calculated across technical replicates within each transect. Dashed lines show linear regression trends across transects, with shaded areas indicating 95% confidence intervals. Adjusted $R^2$ values and p-values from linear models are shown for each group.

### Supplementary Tables

**Table S1. Plant species recorded during the plant community survey.** Summary of the 56 plant species observed along the five transects diversity monitoring over Spring 2024 at the Southern Light Farm, including taxonomic identity and classification by functional role, vertical stratum, and growth form.

| Plant_family | Plant_species | Common_name | Functional_role | Strata | Plant_form |
|---|---|---|---|---|---|
| Apiaceae | *Anthriscus sp.* | Chervil | Spontaneous | Ground cover | Herbaceous |
| Asteraceae | *Artemisia absinthium* | Wormwood | Supportive | Low | Bush |
| Asteraceae | *Bidens bipinnata* | Spanish needle | Supportive | Ground cover | Herbaceous |
| Asteraceae | *Carduus sp.* | Milk thistle | Spontaneous | Ground cover | Herbaceous |
| Asteraceae | *Helminthotheca echioides* | Bristly oxtongue | Spontaneous | Ground cover | Herbaceous |
| Asteraceae | *Taraxacum officinale* | Dandelion | Spontaneous | Ground cover | Herbaceous |
| Asteraceae | *Tithonia diversifolia* | Mexican sunflower | Supportive | Medium | Bush |
| Boraginaceae | *Cynoglossum creticum* | Blue Hound's-tongue | Spontaneous | Ground cover | Herbaceous |
| Brassicaceae | *Sinapis arvensis* | Wild mustard | Spontaneous | Ground cover | Herbaceous |
| Caryophyllaceae | *Stellaria media* | Chickweed | Spontaneous | Ground cover | Herbaceous |
| Commelinaceae | *Tradescantia pallida* | Purple heart | Supportive | Ground cover | Herbaceous |
| Convolvulaceae | *Convolvulus sp.* | Bindweed | Spontaneous | Ground cover | Herbaceous |
| Euphorbiaceae | *Ricinus communis* | Castor oil plant | Supportive | Medium | Bush |
| Fabaceae | *Acacia dealbata* | Mimosa | Supportive | High | Tree |
| Fabaceae | *Erythrostemon gilliesii* | Bird of paradise | Supportive | Medium | Tree |
| Fabaceae | *Medicago polymorpha* | Bur clover | Supportive | Medium | Herbaceous |
| Fabaceae | *Vicia sativa* | Common vetch | Supportive | Medium | Herbaceous |
| Geraniaceae | *Geranium molle* | Dove's-foot crane's-bill | Spontaneous | Ground cover | Herbaceous |
| Juglandaceae | *Juglans regia* | Walnut tree | Supportive | Emergent | Tree |
| Lamiaceae | *Lamium bifidum* | White henbit | Spontaneous | Ground cover | Herbaceous |
| Lamiaceae | *Plectranthus barbatus* | Toilet paper plant | Supportive | Low | Bush |
| Lamiaceae | *Rosmarinus officinalis* | Rosemary | Supportive | Low | Bush |
| Lauraceae | *Laurus nobilis* | Bay laurel | Supportive | Medium | Bush |
| Lythraceae | *Punica granatum* | Pomegranate | Supportive | Medium | Bush |
| Malvaceae | *Gossypium herbaceum* | Cotton plant | Supportive | Low | Bush |
| Malvaceae | *Malva sylvestris* | Common mallow | Spontaneous | Ground cover | Herbaceous |
| Moraceae | *Ficus carica* | Fig tree | Supportive | High | Tree |
| Moraceae | *Morus sp.* | Mulberry tree | Supportive | High | Tree |
| Myrtaceae | *Acca sellowiana* | Feijoa | Supportive | Medium | Bush |
| Myrtaceae | *Eucalyptus globulus* | Eucalyptus tree | Supportive | Emergent | Tree |
| Oleaceae | *Olea europaea* | Olive tree | Crop | High | Tree |
| Oxalidaceae | *Oxalis pes-caprae* | African wood-sorrel | Spontaneous | Ground cover | Herbaceous |
| Oxalidaceae | *Oxalis pes-caprae var. pleniflora* | Cape sorrel | Spontaneous | Ground cover | Herbaceous |
| Papaveraceae | *Fumaria capreolata* | White fumitory | Spontaneous | Ground cover | Herbaceous |
| Papaveraceae | *Fumaria sp.* | Pink fumitory | Spontaneous | Ground cover | Herbaceous |
| Primulaceae | *Lysimachia arvensis* | Red pimpernel | Spontaneous | Ground cover | Herbaceous |
| Primulaceae | *Lysimachia foemina* | Blue pimpernel | Spontaneous | Ground cover | Herbaceous |
| Primulaceae | *Lysimachia foemina v. white* | White pimpernel | Spontaneous | Ground cover | Herbaceous |
| Ranunculaceae | *Ranunculus repens* | Lady's mantle like | Spontaneous | Ground cover | Herbaceous |

| | | | | | |
|---|---|---|---|---|---|
| Rosaceae | *Malus sp.* | Apple tree | Supportive | High | Tree |
| Rosaceae | *Prunus armeniaca* | Apricot tree | Supportive | Medium | Tree |
| Rosaceae | *Prunus persica* | Peach tree | Supportive | Medium | Tree |
| Rosaceae | *Prunus sp.* | Prune tree | Supportive | Medium | Tree |
| Rosaceae | *Rubus ulmifolius* | Wild blackberry | Spontaneous | Medium | Bush |
| Rubiaceae | *Galium aparine* | Cleavers | Spontaneous | Ground cover | Herbaceous |
| Rutaceae | *Citrus aurantiifolia* | Lime tree | Crop | Medium | Tree |
| Rutaceae | *Citrus limon* | Lemon tree | Crop | Medium | Tree |
| Rutaceae | *Citrus reticulata* | Mandarin tree | Crop | Medium | Tree |
| Rutaceae | *Citrus sinensis* | Orange tree Valencia | Crop | Medium | Tree |
| Rutaceae | *Citrus sp.* | Citrus tree | Crop | Medium | Tree |
| Salicaceae | *Populus alba* | White poplar | Supportive | Emergent | Tree |
| Scrophulariaceae | *Scrophularia peregrina* | Mediterranean figwort | Spontaneous | Ground cover | Herbaceous |
| Solanaceae | *Solanum muricatum* | Pepino melon | Supportive | Low | Bush |
| Solanaceae | *Solanum nigrum* | Black nightshade | Spontaneous | Ground cover | Herbaceous |
| Urticaceae | *Parietaria judaica* | Pellitory of the wall | Spontaneous | Ground cover | Herbaceous |
| Urticaceae | *Urtica dioica* | Stinging nettle | Spontaneous | Ground cover | Herbaceous |

**Table S2. Insect taxa recorded during transect surveys.** Summary of the 43 insect species (34 families) recorded across all transects during Spring 2024, identified to species or genus when possible and otherwise grouped into operational morphogroups. Taxa are classified by morphogroup, taxonomic identity, common name, and assigned ecological role in relation to crop production.

| Insect_ morphogroup | Insect_family | Insect_species | Insect_common_name | Ecological_role |
|---|---|---|---|---|
| Antlions | Nemopteridae | *Nemoptera coa* | Greek Spoonwing | Limited role in crops |
| Ants | Formicidae | *Camponotus sp.* | Carpenter ant | Mixed |
| Ants | Formicidae | *Crematogaster schmidti* | Schmidt's Cocktail Ant | Mixed |
| Ants | Formicidae | *Lasius sp.* | Black garden ant | Mixed |
| Beetles | Curculionidae | *Malvaevora timida* | Timid Mallow Weevil | Potential pests |
| Beetles | Glaphyridae | *Eulasia pareyssei* | | Beneficial pollinators |
| Beetles | Kateretidae | *Brachypterus sp.* | Flower beetle | Beneficial pollinators |
| Beetles | Meloidae | *Mylabris sp.* | Northern Bisterbeetle | Mixed |
| Beetles | Mordellidae | *Mordellini sp.* | Tumbling Flower Beetles | Beneficial pollinators |
| Beetles | Scarabaeidae | *Omaloplia sp.* | | Mixed |
| Beetles | Scarabaeidae | *Oxythyrea funesta* | White Spotted Rose Chafer beetle | Mixed |
| Beetles | Scarabaeidae | *Protaetia cuprea* | Copper chafer | Mixed |
| Beetles | Scarabaeidae | *Tropinata sp.* | | Potential pests |
| Beetles | Tenebrionidae | *Omophlus sp.* | Darkling Beetles | Potential pests |
| Bumblebees | Apidae | *Bombus terrestris* | Bumble bee | Beneficial pollinators |
| Butterflies | Nymphalidae | *Pararge aegeria* | Speckled Wood | Beneficial pollinators |
| Butterflies | Nymphalidae | *Vanessa atalanta* | Red Admiral | Beneficial pollinators |
| Butterflies | Nymphalidae | *Vanessa cardui* | Painted Lady | Beneficial pollinators |
| Butterflies | Pieridae | *Pieris rapae* | Cabbage white | Mixed |
| Crickets | Tettigoniidae | *Eupholidoptera megastyla* | Greek Marbled Bush-Cricket | Potential pests |
| Dragonflies | Coenagrionidae | *Coenagrion sp.* | Narrow-winged Damselflies | Natural predators |
| Dragonflies | Lestidae | *Lestes barbarus* | Migrant Spreadwing | Natural predators |
| Dragonflies | Libellulidae | *Sympetrum striolaturm* | Common Darter | Natural predators |
| Flies | Asilidae | *Dioctria sp.* | Robber fly | Natural predators |
| Flies | Empididae | *Hilara sp.* | Balloon fly | Natural predators |
| Flies | Sarcophagidae | *Sarcophaga sp.* | Common Flesh Flies | Limited role in crops |
| Grasshoppers | Acrididae | *Aiolopus sp.* | Green-winged Grasshoppers | Potential pests |
| Grasshoppers | Acrididae | *Anacridium aegyptium* | Egyptian Bird Grasshopper | Potential pests |
| Honeybees | Apidae | *Apis mellifera* | Honeybee | Beneficial pollinators |
| Hoverflies | Syrphidae | *Episyrphus balteatus* | Marmalade Hoverfly | Mixed |
| Hoverflies | Syrphidae | *Xanthogramma sp.* | Harlequin Flies | Mixed |
| Large bees | Andrenidae | *Andrena sp.* | Mining bees | Beneficial pollinators |
| Large bees | Apidae | *Eucera sp.* | Long-horned bees | Beneficial pollinators |
| Large bees | Apidae | *Xylocapa violacea* | Carpenter bee | Beneficial pollinators |
| Moths | Noctuidae | *Acontia trabealis* | Spotted Sulphur | Mixed |
| Scorpionflies | Panorpidae | *Panorpa lacedaemonia* | Scorpionfly | Limited role in crops |
| Small bees | Halictidae | *Lasioglossum sp.* | Sweat bee | Beneficial pollinators |
| True bugs | Cydnidae | *Cydnus aterrimus* | Black Burrowing Bug | Potential pests |
| True bugs | Miridae | *Closterotomus sp.* | | Potential pests |
| True bugs | Miridae | *Deraeocoris ruber* | Red-spotted Plant Bug | Natural predators |
| True bugs | Pentatomidae | *Graphosoma italicum* | Striped Shield bug | Potential pests |

| | | | | |
|---|---|---|---|---|
| True bugs | Rhyparochromidae | *Beosus quadripunctatus* | Dirt-colored seed bug | Potential pests |
| Wasps | Chrysididae | *Holopyga fervida* | Cuckoo wasp | Natural predators |

**Table S3. List of the soil fungal genera detected through metagenomic across the farm.** The 25 fungal genera (18 families) and their respective mean relative abundance (mean_pct), standard deviation (sd_pct), and abundance range (range_pct) across all transects during Spring 2024 are shown. Genus functional lifestyles were assigned based on the FungalTraits database (Põlme et al., 2020). Plant pathogenic (including plant_pathogen and some foliar_endophyte) are fungi infecting living plant tissues and deriving nutrients from host damage; this group includes necrotrophs, hemibiotrophs, and latent pathogens. Saprotrophic (including litter_saprotroph, soil_saprotroph, wood_saprotroph, and unspecified_saprotroph) are fungi that decompose dead organic matter to obtain nutrients and are key drivers of carbon and nutrient cycling. Symbiotic (including arbuscular_mycorrhizal and root-associated) are mutualistic or commensal fungi associated with plant tissues that may enhance plant performance. Animal-associated (including animal_parasite, animal_decomposer, and mycoparasite) are fungi that parasitise or decompose animals (e.g., entomopathogens or nematophagous fungi). Predicted species correspond to the most abundant species-level taxonomic assignment.

| Family | Genus | Lifestyles | mean – pct | sd_ pct | range_pct | Predicted_ species |
|---|---|---|---|---|---|---|
| Nectriaceae | Fusarium | plant_pathogen / litter_saprotroph | 31,19 | 8,82 | 21.55 - 46.01 | *F. flagelliforme** |
| Aspergillaceae | Aspergillus | unspecified_saprotroph / foliar_endophyte | 20,81 | 13,4 | 4.9 - 44.28 | *A. versicolor* |
| Glomeraceae | Rhizophagus | arbuscular_mycorrhizal / root-associated | 15,46 | 15,09 | 4.04 - 45.51 | *R. irregularis* |
| Aspergillaceae | Penicillium | unspecified_saprotroph / foliar_endophyte | 7,25 | 1,72 | 3.19 - 8.98 | *P. citrinum.* |
| Hypocreaceae | Trichoderma | mycoparasite / foliar_endophyte | 4,17 | 1,42 | 2.24 - 6.84 | *T. gamsii; T. atroviride; T. simmonsii* |
| Clavicipitaceae | Metarhizium | animal_parasite / animal_decomposer | 2,98 | 3,97 | 0.18 - 11.88 | *M. brunneum* |
| Pleosporaceae | Alternaria | plant_pathogen / litter_saprotroph | 1,67 | 0,99 | 0.45 - 3.39 | *A. atra** |
| Bionectriaceae | Clonostachys | wood_saprotroph / plant_pathogen | 1,55 | 0,61 | 0.41 - 2.64 | *C. rosea* |
| Ophiocordycipitaceae | Purpureocillium | animal_parasite / animal_decomposer | 1,52 | 0,63 | 0.62 - 3.02 | *P. lilacinum* |
| Trichocomaceae | Talaromyces | unspecified_saprotroph | 1,46 | 0,91 | 0.46 - 3.33 | *T. atroroseus* |
| Glomerellaceae | Colletotrichum | plant_pathogen / litter_saprotroph | 1,28 | 0,35 | 0.76 - 2.13 | *C. gigasporum*** |
| Didymellaceae | Stagonosporopsis | plant_pathogen | 1,16 | 0,88 | 0.22 - 3.01 | *S. rhizophila* |
| Dacampiaceae | Aaosphaeria | soil_saprotroph | 1,16 | 1,29 | 0.18 - 4.58 | *A. arxii* |
| Malasseziaceae | Malassezia | soil_saprotroph / root-associated | 1,12 | 0,37 | 0.47 - 1.78 | *M. furfur* |
| Aspergillaceae | Monascus | unspecified_saprotroph | 1,05 | 2,09 | 0 - 5.97 | *M. purpureus* |
| Apiosporaceae | Nigrospora | litter_saprotroph | 0,98 | 0,45 | 0.44 - 1.97 | *N. osmanthi* |
| Chaetomiaceae | Thermothielavioides |  | 0,73 | 0,25 | 0.37 - 1.39 | *T. terrestris* |
| Chaetomiaceae | Thermothelomyces | soil_saprotroph | 0,72 | 0,33 | 0.15 - 1.68 | *T. thermophilus* |
| Sporidiobolaceae | Rhodotorula | unspecified_saprotroph / foliar_endophyte | 0,72 | 0,23 | 0.21 - 1.21 | *R. graminis* |
| Didymellaceae | Boeremia | plant_pathogen / foliar_endophyte | 0,63 | 0,48 | 0 - 1.78 | *B. exigua* |
| Chaetomiaceae | Chaetomium | litter_saprotroph / foliar_endophyte | 0,62 | 0,29 | 0.14 - 1.2 | *C. globosum* |
| Didymellaceae | Ascochyta | plant_pathogen | 0,62 | 0,25 | 0.25 - 1.21 | *A. rabiei* |
| Didymellaceae | Macroventuria | litter_saprotroph | 0,6 | 0,44 | 0.18 - 1.62 | *M. anomochaeta* |
| Cucurbitariaceae | Cucurbitaria | plant_pathogen / litter_saprotroph | 0,35 | 0,45 | 0.03 - 1.39 | *C. berberidis* |
| Wallemiaceae | Wallemia | unspecified_saprotroph | 0,21 | 0,31 | 0 - 0.93 | *W. mellicola* |

** Lab and field verified species

* Visual field unverified species

**Table S4. List of the plant genera detected through soil metagenomic across the farm.** The 29 plant genera (14 families) and their respective mean relative abundance (mean_pct), standard deviation (sd_pct), and abundance range (range_pct) across all transects during Spring 2024 are shown. Metagenomic plant genera were validated and categorised (Field_verif) using on-site transect surveys, farmer-reported planting and management history, and regional occurrence records. Predicted species correspond to the most abundant species-level taxonomic assignment.

| Family | Genus | Plant_form | Field_verif | mean _pct | sd_ pct | range_pct | Predicted_ species |
|---|---|---|---|---|---|---|---|
| Urticaceae | Parietaria | Herbaceous | Direct field verified | 19,52 | 17,38 | 3 - 47.52 | *P. judaica*** |
| Rutaceae | Citrus | Tree | Direct field verified | 8,63 | 5,59 | 1.4 - 16 | *C. sinensis**, C. reticulata**, C. aurantiifolia*,C. limon** |
| Oleaceae | Olea | Tree | Direct field verified | 8,35 | 7,95 | 1.45 - 21.33 | *O. europaea*** |
| Fabaceae | Vicia | Herbaceous | Direct field verified | 7,98 | 15,77 | 0 - 36.12 | *V. sativa*** |
| Urticaceae | Urtica | Herbaceous | Direct field verified | 6,72 | 5,42 | 0.48 - 13.16 | *U. dioica** |
| Euphorbiaceae | Mercurialis | Herbaceous | Regionally plausible | 5,91 | 9,99 | 0.4 - 23.72 | *M. annua* |
| Oleaceae | Fraxinus | Tree | Regionally plausible | 5,82 | 4,99 | 1.77 - 13.97 | *F. pennsylvanica.(F. ornus)* |
| Poaceae | Hordeum | Herbaceous | Crop/support vegetation | 5,05 | 6,64 | 0.1 - 16.71 | *H. vulgare* |
| Poaceae | Avena | Herbaceous | Crop/support vegetation | 3,58 | 7,18 | 0.11 - 16.41 | *A. sativa* |
| Moraceae | Morus | Tree | Direct field verified | 3,52 | 4,63 | 0.01 - 11.57 | *M. notabilis. (M. nigra)* |
| Poaceae | Brachypodium | Herbaceous | Regionally plausible | 3,05 | 3,56 | 0.2 - 9.24 | *B. distachyon* |
| Poaceae | Secale | Herbaceous | Crop/support vegetation | 2,37 | 2,98 | 0.01 - 7.4 | *S. cereale* |
| Poaceae | Triticum | Herbaceous | Crop/support vegetation | 2,01 | 2,71 | 0.22 - 6.8 | *T. aestivum* |
| Fabaceae | Medicago | Herbaceous | Direct field verified | 1,72 | 2,09 | 0.25 - 5.22 | *M. polymorpha** |
| Fabaceae | Trifolium | Herbaceous | Regionally plausible | 1,7 | 1,64 | 0.59 - 4.56 | *T. repens* |
| Poaceae | Setaria | Herbaceous | Regionally plausible | 1,6 | 1 | 0.94 - 3.3 | *S. viridis* |
| Fabaceae | Pisum | Herbaceous | Crop/support vegetation | 1,47 | 2,84 | 0 - 6.53 | *P. sativum* |
| Solanaceae | Solanum | Bush / Herbaceous | Direct field verified | 1,36 | 0,82 | 0.56 - 2.74 | *S. muricatum**, S. nigrum*** |
| Poaceae | Oryza | Herbaceous | Unlikely / false positive | 1,32 | 0,65 | 0.44 - 2.1 | |
| Mamiellaceae | Micromonas | Herbaceous | Unlikely / false positive | 1,27 | 0,76 | 0.38 - 2.38 | |
| Poaceae | Digitaria | Herbaceous | Regionally plausible | 1,24 | 0,63 | 0.49 - 2.2 | *D. exilis. (D. sanguinalis)* |
| Poaceae | Phragmites | Herbaceous | Regionally plausible | 1,23 | 0,69 | 0.45 - 2.11 | *P. australis* |
| Chlamydomonadaceae | Chlamydomonas | Other | Unlikely / false positive | 0,97 | 0,56 | 0.35 - 1.82 | |
| Chlorellaceae | Chlorella | Other | Unlikely / false positive | 0,82 | 0,47 | 0.21 - 1.47 | *C. sorokiniana* |
| Simaroubaceae | Ailanthus | Tree | Regionally plausible | 0,64 | 0,28 | 0.42 - 1.11 | *A. altissimus* |
| Caryophyllaceae | Agrostemma | Herbaceous | Regionally plausible | 0,61 | 0,42 | 0.19 - 1.3 | *A. githago* |
| Malvaceae | Gossypium | Bush | Direct field verified | 0,61 | 0,28 | 0.43 - 1.1 | *G. herbaceum*** |
| Fagaceae | Quercus | Tree | Regionally plausible | 0,54 | 0,32 | 0.28 - 1.08 | *Q. variabilis. (Q. coccifera)* |
| Poaceae | Bromus | Herbaceous | Regionally plausible | 0,43 | 0,61 | 0.01 - 1.5 | *B. tectorum* |

** Field verified species

* Unverified species but regionally found

. Likely wrong species assignment (most probable species found in the region)

**Table S5. List of the invertebrate genera detected through soil metagenomic across the farm.** The 23 invertebrate genera (19 families) and their respective mean relative abundance (mean_pct), standard deviation (sd_pct), and abundance range (range_pct) across all transects during Spring 2024 are shown Metagenomic invertebrate genera were validated and categorised (Field_verif) using on-site transect surveys, farmer-reported observations and regional occurrence records. Predicted species correspond to the most abundant species-level taxonomic assignment.

| **Family** | **Genus** | **Field_verif** | **mean_ pct** | **sd_ pct** | **range_pct** | **Predicted_Species** |
|---|---|---|---|---|---|---|
| Drosophilidae | Drosophila | Regionally plausible | 11,54 | 2,78 | 7.56 - 16.61 | *D. funebris. (D. subobscura)* |
| Rhabditidae | Oscheius | Unlikely / false positive | 10,28 | 13,2 | 0 - 34.31 | *O. tipulae* |
| Culicidae | Anopheles | Regionally plausible | 6,86 | 1,59 | 3.7 - 10.33 | *A. stephensi. (A. maculipennis)* |
| Limnephilidae | Limnephilus | Regionally plausible | 6,45 | 2,33 | 3.35 - 12.19 | *L. marmoratus. (L. lunatus)* |
| Pollicipedidae | Capitulum | Unlikely / false positive | 5,8 | 1,67 | 2.94 - 9.32 | *C.mitella* |
| Geometridae | Eupithecia | Regionally plausible | 5,2 | 1,95 | 1.96 - 9.09 | *E. insigniata** |
| Timematidae | Timema | Unlikely / false positive | 4,97 | 1,67 | 2.48 - 7.86 | *T. cristinae* |
| Tortricidae | Acleris | Regionally plausible | 3,95 | 1,47 | 2.16 - 7.96 | *Acleris cristana** |
| Formicidae | Lasius | Direct field verified | 3,95 | 4,28 | 0 - 10.06 | *L. fuliginosus** |
| Coelopidae | Coelopa | Regionally plausible | 3,76 | 0,85 | 2.37 - 5.43 | *C.pilipes. (C.Frigida)* |
| Cytherideidae | Cyprideis | Unlikely / false positive | 3,69 | 1,49 | 1.08 - 6.47 | *C. torosa* |
| Noctuidae | Mythimna | Regionally plausible | 3,43 | 1,42 | 1.28 - 6.16 | *M. l-album** |
| Tortricidae | Epinotia | Regionally plausible | 3,39 | 1,04 | 1.5 - 5.11 | *E.demarniana** |
| Erebidae | Eilema | Regionally plausible | 3,36 | 1,29 | 1.69 - 5.9 | *E. caniola** |
| Kateretidae | Brachypterus | Direct field verified | 3,19 | 5 | 0 - 13.53 | *B. glaber* |
| Noctuidae | Xestia | Regionally plausible | 3,01 | 1,46 | 0.85 - 5.74 | *X. c-nigrum** |
| Coleophoridae | Coleophora | Regionally plausible | 2,75 | 1,47 | 1.22 - 7.28 | *C. flavipennella* |
| Palaemonidae | Macrobrachium | Unlikely / false positive | 2,61 | 1,07 | 0.96 - 4.34 | *M. nipponense* |
| Geometridae | Idaea | Regionally plausible | 2,46 | 1,1 | 1.21 - 5.42 | *I. dimidiata** |
| Ypsolophidae | Ypsolopha | Regionally plausible | 2,36 | 1,05 | 0.78 - 4.83 | *Y. scabrella. (Y. sequella)* |
| Yponomeutidae | Yponomeuta | Regionally plausible | 2,34 | 1,19 | 0.9 - 4.79 | *Y. sedellus* |
| Tortricidae | Apotomis | Unlikely / false positive | 2,33 | 1,13 | 0.69 - 5.15 | *A. betuletana* |
| Nemouridae | Nemurella | Unlikely / false positive | 2,33 | 1,06 | 0.91 - 4.35 | *N. pictetii* |

** Field verified species

* Unverified species but regionally found

. Likely wrong species assignment (most probable species found in the region)

Table S6. Genera showing significant differential abundance following agroforestry transition. List of the 38 genera (27 families) whose relative abundance differed significantly between agroforestry transition transects (A–D) and the unconverted reference transect (E). Differential abundance was tested at the genus level from the Kingdoms Fungi, Plantae, and Invertebrate (Arthropoda, Nematoda, and Mollusca phyla only) detected through soil metagenomic analysis using ANCOM-BC2. Only genera passing the FDR threshold ($q < 0.05$) and showing at least a two-fold change relative to the reference transect ($|log2FC| \geq 1$) are reported. Contrast indicates the agroforestry transect compared to E. q is the Benjamini–Hochberg adjusted p-value, and log2FC represents the $log_2$ fold change in abundance relative to E (positive values indicate enrichment, negative values depletion).

| **Contrast** | **Kingdom** | **Family** | **Genus** | **q** | **log2FC** |
|---|---|---|---|---|---|
| A | Fungi | Clavicipitaceae | Metarhizium | 2,99E-09 | -4,50866 |
| A | Fungi | Cucurbitariaceae | Cucurbitaria | 5,84E-08 | 3,089264 |
| A | Fungi | Glomeraceae | Rhizophagus | 1,2E-07 | 1,626005 |
| A | Fungi | Dacampiaceae | Aaosphaeria | 4,23E-06 | -2,3134 |
| A | Fungi | Pleosporaceae | Alternaria | 6,06E-06 | 1,847033 |
| A | Fungi | Ophiocordycipitaceae | Purpureocillium | 1,98E-05 | 1,303712 |
| A | Fungi | Trichocomaceae | Talaromyces | 3,82E-05 | -1,88499 |
| A | Fungi | Bionectriaceae | Clonostachys | 4,93E-05 | -1,61321 |
| A | Plantae | Oleaceae | Olea | 2,46E-11 | 5,388131 |
| A | Plantae | Oleaceae | Fraxinus | 5,21E-11 | 4,438156 |
| A | Plantae | Urticaceae | Urtica | 2,41E-10 | 2,404209 |
| A | Plantae | Rutaceae | Citrus | 5,19E-09 | 1,726199 |
| A | Plantae | Fabaceae | Medicago | 4,65E-08 | 4,791318 |
| A | Plantae | Poaceae | Hordeum | 1,29E-07 | 2,231122 |
| A | Plantae | Poaceae | Brachypodium | 1,11E-06 | -2,18336 |
| A | Plantae | Poaceae | Triticum | 2,71E-05 | 2,783485 |
| A | Animalia | Rhabditidae | Oscheius | 5,84E-07 | 5,273052 |
| A | Animalia | Kateretidae | Brachypterus | 0,00335 | 4,257161 |
| A | Animalia | Noctuidae | Xestia | 0,007342 | -1,03401 |
| B | Fungi | Clavicipitaceae | Metarhizium | 7,27E-10 | -3,26581 |
| B | Fungi | Pleosporaceae | Alternaria | 4,21E-08 | 2,67926 |
| B | Fungi | Dacampiaceae | Aaosphaeria | 1,63E-07 | -2,38558 |
| B | Fungi | Didymellaceae | Stagonosporopsis | 2,21E-06 | 1,958171 |
| B | Fungi | Didymellaceae | Macroventuria | 8,58E-06 | 2,060614 |
| B | Fungi | Didymellaceae | Ascochyta | 0,000115 | 1,109079 |
| B | Fungi | Trichocomaceae | Talaromyces | 0,000449 | -1,35326 |
| B | Fungi | Wallemiaceae | Wallemia | 0,009287 | 3,726444 |
| B | Plantae | Poaceae | Avena | 1,95E-13 | 5,568938 |
| B | Plantae | Poaceae | Hordeum | 2,76E-10 | 3,797859 |
| B | Plantae | Oleaceae | Olea | 5,18E-09 | 3,438201 |
| B | Plantae | Urticaceae | Parietaria | 1,19E-08 | -1,89365 |
| B | Plantae | Oleaceae | Fraxinus | 4,21E-08 | 2,603149 |
| B | Plantae | Poaceae | Triticum | 1,59E-07 | 4,37786 |
| B | Plantae | Euphorbiaceae | Mercurialis | 0,000132 | 1,520256 |
| B | Animalia | Formicidae | Lasius | 1,93E-05 | 4,323815 |
| B | Animalia | Rhabditidae | Oscheius | 4,41E-05 | 3,688747 |
| C | Fungi | Clavicipitaceae | Metarhizium | 2,65E-09 | -4,88776 |
| C | Fungi | Pleosporaceae | Alternaria | 6,95E-07 | 2,388961 |
| C | Fungi | Aspergillaceae | Aspergillus | 1,01E-06 | 1,253425 |

| | | | | | |
|---|---|---|---|---|---|
| C | Fungi | Dacampiaceae | Aaosphaeria | 1,82E-05 | -3,02468 |
| C | Fungi | Hypocreaceae | Trichoderma | 5,38E-05 | 1,265271 |
| C | Fungi | Trichocomaceae | Talaromyces | 0,000259 | -1,35503 |
| C | Plantae | Euphorbiaceae | Mercurialis | 6,06E-10 | 4,920025 |
| C | Plantae | Urticaceae | Urtica | 1,86E-08 | 2,07828 |
| C | Plantae | Rutaceae | Citrus | 1,04E-06 | -1,83809 |
| C | Plantae | Poaceae | Phragmites | 0,000138 | 1,22823 |
| C | Plantae | Oleaceae | Olea | 0,000165 | 1,650294 |
| C | Plantae | Poaceae | Hordeum | 0,000195 | 1,440957 |
| C | Plantae | Poaceae | Triticum | 0,007387 | 1,777243 |
| C | Animalia | Formicidae | Lasius | 6,27E-05 | 4,024966 |
| C | Animalia | Rhabditidae | Oscheius | 0,335646 | -1,09004 |
| D | Fungi | Aspergillaceae | Aspergillus | 1,13E-09 | -2,77692 |
| D | Fungi | Glomeraceae | Rhizophagus | 1,88E-08 | 2,299102 |
| D | Fungi | Clavicipitaceae | Metarhizium | 3,53E-07 | -2,44487 |
| D | Fungi | Dacampiaceae | Aaosphaeria | 4,13E-05 | -3,11864 |
| D | Fungi | Didymellaceae | Stagonosporopsis | 7,7E-05 | -1,87428 |
| D | Fungi | Aspergillaceae | Penicillium | 7,7E-05 | -1,57643 |
| D | Fungi | Bionectriaceae | Clonostachys | 0,000545 | -1,35059 |
| D | Fungi | Chaetomiaceae | Thermothielavioides | 0,001222 | -1,43083 |
| D | Fungi | Chaetomiaceae | Chaetomium | 0,003401 | -2,08928 |
| D | Fungi | Trichocomaceae | Talaromyces | 0,009449 | -1,06516 |
| D | Fungi | Chaetomiaceae | Thermothelomyces | 0,010896 | -1,8141 |
| D | Plantae | Oleaceae | Olea | 3,77E-10 | 4,809176 |
| D | Plantae | Rutaceae | Citrus | 3,77E-10 | 3,167192 |
| D | Plantae | Poaceae | Brachypodium | 1,13E-09 | 4,310388 |
| D | Plantae | Oleaceae | Fraxinus | 1,6E-09 | 3,905313 |
| D | Plantae | Poaceae | Setaria | 9,88E-08 | 3,405031 |
| D | Plantae | Fabaceae | Trifolium | 1,64E-07 | 4,057003 |
| D | Plantae | Fabaceae | Medicago | 2,21E-07 | 4,418655 |
| D | Plantae | Urticaceae | Parietaria | 5,53E-06 | -1,60279 |
| D | Plantae | Euphorbiaceae | Mercurialis | 1,61E-05 | 2,229153 |
| D | Plantae | Poaceae | Hordeum | 0,0001 | -1,98787 |
| D | Plantae | Urticaceae | Urtica | 0,002012 | -1,2739 |
| D | Plantae | Malvaceae | Gossypium | 0,006006 | 1,134014 |
| D | Animalia | Nemouridae | Nemurella | 0,023548 | -1,15911 |

Table S7. Summary statistics of metagenomic assembly and functional annotation across soil samples and at the farm level.

| **Metric** | **A** | **B** | **C** | **D** | **E** | **Farm** |
|---|---|---|---|---|---|---|
| Nb of contig | 84.906 | 42.755 | 34.349 | 6.561 | 43.520 | **212.091** |
| Nb of predicted protein | 102.872 | 51.561 | 41.065 | 7.719 | 54.534 | **257.751** |
| Protein average length (pb) | 477,98 | 499,26 | 457,80 | 417,18 | 568,70 | **496,39** |
| Protein min length (pb) | 200 | 207 | 200 | 229 | 200 | **200** |
| Protein max length (pb) | 3.164 | 5.338 | 2.224 | 3.506 | 5.754 | **5.754** |
| Protein SD length (pb) | 203,44 | 283,28 | 166,54 | 173,91 | 372,89 | **262,64** |
| Nb of protein taxonomically classified | 34.320 | 16.280 | 14.761 | 2.794 | 13.899 | **82.054** |
| Nb of bacterial protein (Kingdom) | 28.918 | 13.052 | 12.477 | 2.306 | 10.703 | **67.456** |
| Nb of bacterial protein (Class) | 8.169 | 4.480 | 3.762 | 403 | 2.329 | **19.143** |
| Nb of bacterial protein (Genus) | 6.153 | 3.633 | 3.038 | 311 | 1.920 | **15.055** |
| Nb of unique Genus | 220 | 197 | 146 | 59 | 123 | **346** |
| Nb of unique protein family (> 3 protein members) | 524 | 126 | 143 | 87 | 191 | **988** |
| Number of unique Pfam | 343 | 102 | 117 | 91 | 165 | **416** |
| Nb of protein family with Pfam | 467 | 113 | 131 | 79 | 175 | **889** |
| Nb of unique Gene Ontology (GO) | 50 | 30 | 27 | 27 | 37 | **55** |
| Nb of unique functional units (Pfam-GO) | 205 | 72 | 98 | 76 | 125 | **253** |
| Nb of protein without functional annotation | 57 | 13 | 12 | 8 | 16 | **99** |

Table S8. Outcome of the enrichment analysis of nutrient-related Pfam domains across agroforestry transects. All 18 Pfam domains retained after manual curation (see Method) are reported. Contrast indicates the agroforestry transect compared to the reference transect E. Relative abundances of Pfam-associated protein families were compared between agroforestry transects (A–D) and the unconverted reference transect (E) using Fisher's exact tests. q-values correspond to FDR-adjusted p-values. Log2FC represents the $log_2$ fold change in abundance relative to E (positive values indicate enrichment; negative values indicate depletion). "Inf" in Log2FC values reflect the absence of the Pfam domain in the reference transect, indicating a absolute gain or lost of function.

| Nutrient category | Contrast | Pfam hit | Pfam ID | Total count | q | Log2FC |
|---|---|---|---|---|---|---|
| Carbohydrate | A | PF00083 | Sugar_tr | 48 | 9,97E-07 | Inf |
| Carbohydrate | A | PF00128 | Alpha-amylase | 108 | 1,94E-15 | Inf |
| Carbohydrate | A | PF00591 | Glycos_transf_3 | 9 | 0,185298 | Inf |
| Carbohydrate | A | PF00723 | Glyco_hydro_15 | 6 | 0,600289 | -1,28 |
| Carbohydrate | A | PF00982 | Glyco_transf_20 | 30 | 0,000265 | Inf |
| Carbohydrate | A | PF01380 | SIS | 18 | 0,012257 | Inf |
| Carbohydrate | B | PF00343 | Phosphorylase | 36 | 3,94E-13 | Inf |
| Carbohydrate | B | PF00723 | Glyco_hydro_15 | 6 | 0,73185 | -Inf |
| Carbohydrate | C | PF00723 | Glyco_hydro_15 | 6 | 0,705483 | -Inf |
| Carbohydrate | C | PF00953 | Glycos_transf_4 | 21 | 1,55E-06 | Inf |
| Carbohydrate | D | PF00723 | Glyco_hydro_15 | 6 | 1 | -Inf |
| Nitrogen | A | PF00120 | Gln-synt_C | 16 | 0,005348 | -2,87 |
| Nitrogen | A | PF00909 | Ammonium_transp | 21 | 0,004625 | Inf |
| Nitrogen | A | PF01645 | Glu_synthase | 12 | 0,393494 | Inf |
| Nitrogen | B | PF00120 | Gln-synt_C | 16 | 0,077445 | -Inf |
| Nitrogen | B | PF01593 | Amino_oxidase | 27 | 8,58E-10 | Inf |
| Nitrogen | B | PF01645 | Glu_synthase | 12 | 0,286941 | Inf |
| Nitrogen | C | PF00120 | Gln-synt_C | 16 | 0,027279 | -Inf |
| Nitrogen | C | PF01645 | Glu_synthase | 12 | 0,423136 | Inf |
| Nitrogen | D | PF00120 | Gln-synt_C | 16 | 1 | 0,78 |
| Nitrogen; Sulfur | A | PF03460 | NIR_SIR_ferr | 27 | 0,000676 | Inf |
| Phosphorus | A | PF01384 | PHO4 | 21 | 0,004625 | Inf |
| Phosphorus | A | PF01658 | Inos-1-P_synth | 13 | 0,393494 | Inf |
| Phosphorus | A | PF03030 | H_PPase | 80 | 5,15E-08 | Inf |
| Phosphorus | B | PF01658 | Inos-1-P_synth | 13 | 0,286941 | Inf |
| Phosphorus | C | PF01658 | Inos-1-P_synth | 13 | 0,263702 | Inf |
| Phosphorus | C | PF03030 | H_PPase | 80 | 1,75E-07 | Inf |
| Sulfur | A | PF00884 | Sulfatase | 12 | 0,393494 | Inf |
| Sulfur | A | PF01507 | PAPS_reduct | 30 | 0,000265 | Inf |
| Sulfur | B | PF00884 | Sulfatase | 12 | 0,286941 | Inf |
| Sulfur | C | PF00884 | Sulfatase | 12 | 0,423136 | Inf |
